# The genetic architecture of cortical similarity networks

**DOI:** 10.1101/2024.12.18.629155

**Authors:** Isaac Sebenius, Varun Warrier, Richard A.I. Bethlehem, Richard Dear, Eva-Maria Stauffer, Yuanjun Gu, Rafael Romero-Garcia, Jakob Seidlitz, Edward Bullmore, Sarah Morgan

**Author notes:** These authors contributed equally to this work (Edward Bullmore, Sarah Morgan).

## Abstract

The genetic architecture of human brain networks is central to understanding the organization and evolution of the cortex, the causal relationships between brain structure and function, and the pathogenesis of heritable neuropsychiatric disorders. However, the current understanding of the genetics of brain networks remains fragmented. Here, we investigated common genetic effects on Morphometric INverse Divergence (MIND), a biologically-validated, heritable, and multimodal MRI metric of inter-areal similarity and connectivity. Using a discovery dataset (N***>***30,000 adults), we estimated subject-specific MIND networks from the multivariate distributions of four MRI features at each of 23 cortical areas and performed genome-wide association studies (GWAS) at each of the 276 inter-areal edges. These edge-level genetic effects were highly replicated by parallel GWAS of an independent validation dataset (N***>***18,000 adults). We found that strong genetic correlations between multiple edges were largely reducible to two gradients of genetically-determined cortical similarity, each of which was aligned with geodesic distance from one of the two phylogenetically primitive areas (paleocortex and archicortex) predicted by the dual origin theory of cortical evolution. Genetic MIND gradients were more heritable than comparable gradients derived from GWAS of functional MRI connectivity networks; and the paleocortical trend was genetically correlated with, and causally predictive of, functional connectivity. Finally, we identified multiple global and local genetic correlations between both MIND gradients and nine clinical diagnoses or biomedical traits, indicating that the normative genetic architecture of human brain networks is pleiotropically associated with inherited risk of neuropsychiatric disorders and systemic metabolic and immune traits. These results provide fresh insight into the dual origins of the human cortex and their implications for brain function and health.

## Introduction

Identifying the genetic basis of the human brain’s complex network structure (Bullmore and Sporns, 2009; Bassett and Sporns, 2017) is essential for understanding how the brain develops, functions, and relates to health and cognition (Dennis et al, 2019). Over a decade of work has shown that individual differences in brain network organization measured by multiple modalities of magnetic resonance imaging (MRI) are under genetic influence (Schutte et al, 2013; Chiang et al, 2009; Glahn et al, 2010; Dennis et al, 2019). Recent genome-wide association studies (GWAS) have begun to investigate the common genetic variation associated with certain brain network phenotypes, including (f)MRI-derived functional connectivity (FC) from correlational analysis of regional time-series, and diffusion (d)MRI-derived structural connectivity (SC) from tractographic reconstruction of axonal projections between regions (Wainberg et al, 2024; Tissink et al, 2023; Roelfs et al, 2024; Foo et al, 2021; Cheng et al, 2024). However, our understanding of genetic influences on brain network organization remains fragmented. For example, it has proven difficult to identify any meaningful genetic overlap between brain structural and functional connectivity phenotypes derived from different MRI modalities (Tissink et al, 2023; Roelfs et al, 2024; Gajwani et al, 2023; Luppi et al, 2024). The genetic architecture of global or localised (regional) MRI measures of brain structure has been well characterized (Smith et al, 2021; Grasby et al, 2020; Zhao et al, 2019; Elliott et al, 2018; Warrier et al, 2023; Fu et al, 2024; Tissink et al, 2024), but it is unclear to what extent these phenotypes are genetically correlated with MRI metrics of network connectivity between localised regions. Most fundamentally, the rapidly growing field of neuroimaging GWAS and post-GWAS studies has been empirically driven, with minimal theoretical frameworks to guide interpretation and integration of results from multi-modal MRI datasets in terms of prior models of the genetic origins and organization of the cortex (Sebenius et al, 2024).

We have begun to address these key knowledge gaps by characterizing the common genetic variation associated with structural MRI similarity, an innovative approach to human cortical network phenotyping (Wang and He, 2024; Sebenius et al, 2024). It has been shown that the structural similarity between cortical areas, measured using one or more macro- and/or micro-structural MRI features, is both heritable and reflective of multiple neurobiological properties, including: the architectonic similarity between areas at a cellular scale (Seidlitz et al, 2018; Paquola et al, 2019; Sebenius et al, 2023); the likelihood and/or strength of inter-areal axonal projections (Beul et al, 2015, 2017; Wei et al, 2018; Seidlitz et al, 2018; Sebenius et al, 2023); and the transcriptomic similarity between areas based on regional bulk whole genome expression (Seidlitz et al, 2018; Sebenius et al, 2023). Here, we used multimodal MRI from N*>*30,000 subjects (and replicated in an additional N*>*18,000) from the UK Biobank (UKB) to estimate GWAS statistics independently for each edge of the structural similarity network estimated using Morphometric INverse Divergence (MIND), a method that has been favourably benchmarked, both technically and biologically, against alternative structural connectivity and similarity estimators (Sebenius et al, 2023). This preliminary analysis demonstrated that MIND network edges have moderate SNP-based heritability (mean 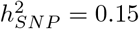) and 54 genomic loci are significantly associated with variation of MIND similarity at one or more of the 276 constituent edges in each individual brain similarity network. On this basis, we addressed three linked questions concerning the genetic architecture of human cortical networks.

The long-standing dual origin theory of cortical evolution and development posits that mammalian cortex evolved along two genetically differentiated cytoarchitectonic trends of expansion and increasing lamination, originating from two specific loci of primitive cortical tissue (Abbie, 1940; Dart, 1934; Sanides, 1971; Pandya et al, 2015; Valk et al, 2020, 2022). If this theory is true, we reasoned, there should be evidence that genetic effects on structural similarity between cortical areas are systematically co-ordinated along two dimensions or gradients originating at or close to the theoretically predicted origins of cortical evolution: the piriform cortex (paleo-cortex) and the hippocampus (archi-cortex). We tested this hypothesis by analyzing the MIND genetic correlation matrix, where each element represents the correlation between the GWAS statistics for each pair of 276 edges in the MIND network. We found that over 70% of the co-variation in the MIND genetic correlation matrix was reducible to two principal gradients, G1 and G2, which were tested for spatial alignment with the two theoretically predicted trends of paleo- and archi-cortical architectonic differentiation (Valk et al, 2020, 2022; Dart, 1934; Sanides, 1971). We found that the weighting of cortical areas on the first genetic gradient was uniquely highly correlated with geodesic distance from the paleo-cortical origin in piriform cortex. Whereas cortical weighting on the second gradient was most correlated with distance from parahippocampal cortex, adjacent to the archi-cortical origin. Building on these results, we investigated two further key questions, through the lens of these dual origin genetic gradients in human MIND networks.

First we investigated the hypothesis that inter-areal metrics of structural similarity and functional connectivity (FC) should be genetically correlated. To that end, we estimated comparable GWAS statistics for single-subject fMRI connectivity networks measured in the same (discovery) cohort of UKB participants, and with identical cortical nodes and edges to the MIND networks. We found, as expected, that FC heritability was significantly less than MIND heritability, overall; but structural similarity across the paleocortical gradient was robustly genetically correlated with functional connectivity. Indeed, for the paleocortical trend specifically, Mendelian randomization analysis indicated that genetically determined variation in structural similarity caused concordant changes in functional connectivity.

Finally, we explored how the genetics of MIND are related to the genetics of clinical disorders or biomedical traits that putatively lie downstream of both structural similarity and functional connectivity phenotypes. Using global and local genetic correlation analysis, we identified multiple genomic loci of pleiotropic association with both dual origin trends of cortical similarity and trans-diagnostic clusters of more than one disorder or trait.

All MIND GWAS summary statistics will be made available to facilitate the replication and extension of these analyses. This open resource includes genome-wide associations both with the two genetic gradients identified here and at the level of individual MIND network edges.

### Genome-wide associations with edges of inter-areal similarity

MIND estimates similarity between cortical areas by comparing their vertex- or voxel-level distributions of one or more structural MRI features. Our aim was to use an integrated measure of structural similarity, representative of multiple, complementary aspects of cortical organization. From the range of MRI features potentially available in the UKB MRI cohort, we therefore selected four MRI features that were individually interpretable and heritable (see Methods for details): two macro-structural metrics - cortical thickness (CT) and mean curvature (MC); and two micro-structural metrics - mean diffusivity (MD) and neurite density index (NDI). For each participant, these four features were measured at every vertex within each of 360 predefined cortical regions (the “HCP” parcellation; Glasser et al (2016)). The similarity between each pair of HCP-defined regional multivariate distributions was estimated by the inverse of the Kullback-Leibler divergence between them (Sebenius et al, 2023). The resulting {360 region × 360 region} networks were coarse-grained by averaging inter-regional similarities for HCP-designated parcels assigned to the same one of 23 higher-order cortical areas. This led to a final set of subject-specific {23 area × 23 area} structural similarity networks, each comprising 276 edges representing MIND similarity between each possible pair of 23 cortical areas (Fig. 1A).

**Fig. 1.**
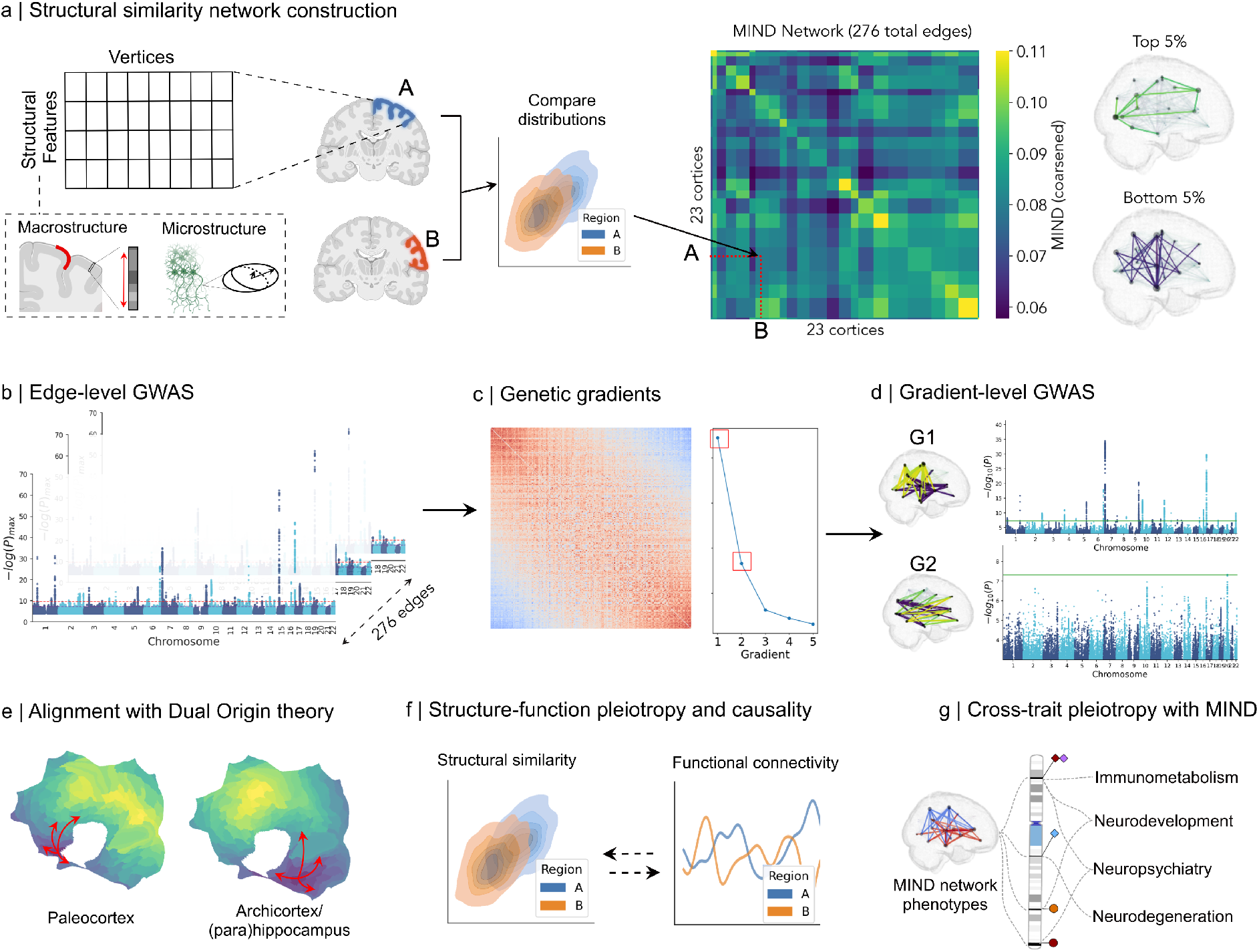
Overview of structural similarity network estimation by MIND and downstream analysis of the dual origins of MIND genetic gradients and their pleiotropic associations with functional and clinical phenotypes. A) Schematic of the process for transforming an individual’s brain MRI scans into a MIND network comprising 276 edges between 27 cortical areas per hemisphere (see **Methods**). The group-averaged MIND network is shown as a heatmap (color bar capped at 0.11 to aid visualization). To the right of the heatmap, and sharing the same colorbar, the top and bottom 5% edges of the MIND network are shown on the brain. B) We performed genome-wide association analyses for each of the 276 MIND network edges, and (C) identified latent genetic gradients underlying the edge-level signal. D) We performed two additional GWAS corresponding to MIND network connectivity summarized along the axes defined by G1 and G2, the primary genetic gradients of MIND. E) We analyzed the spatial variation of the genetically-defined MIND gradients in the context of dual origin theory. F) Using summary statistics from directly-comparable phenotypes from functional connectivity networks, we tested for both correlative and causal genetic overlap between structural similarity and functional connectivity. G) Finally, we examined the global and local genetic overlaps between structural similarity and a variety of traits, including immunometabolic, neurodevelopmental, neuropsychiatric and neurodegenerative phenotypes.

Each of these edges of inter-areal similarity was treated as an independent phenotype in a set of 276 GWAS using a discovery sample of *N*_*max*_=30,524 individuals from the UKB (based on the same primary cohort as in Warrier et al (2023)). MIND edges demonstrated moderate heritability (mean 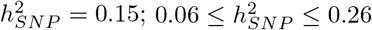). At an experiment-wide threshold of *P* < 1.8 × 10^−10^ (i.e., 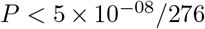, 54 independent genomic loci were associated with at least one of the 276 MIND network edges (Fig. 2A). We replicated these results by performing the same GWAS on an additional and independent validation cohort of *N*_*max*_ =18,769 UKB adult participants. To test generalizability of MIND genetic associations, we computed the genetic correlation using LD Score Regression (LDSC) (Bulik-Sullivan et al, 2015) between each edge in the discovery and validation cohorts (Extended Data Fig. 1). The mean genetic correlation was *r*_*g*_=1.003 (0.57 ≤ *r*_*g*_ ≤ 1.8), with 274 of 276 edges (> 99%) significantly positively correlated (*P*_*FDR*_ < 0.05), indicating robust replicability of the GWAS statistics of MRI similarity network phenotypes.

**Fig. 2.**
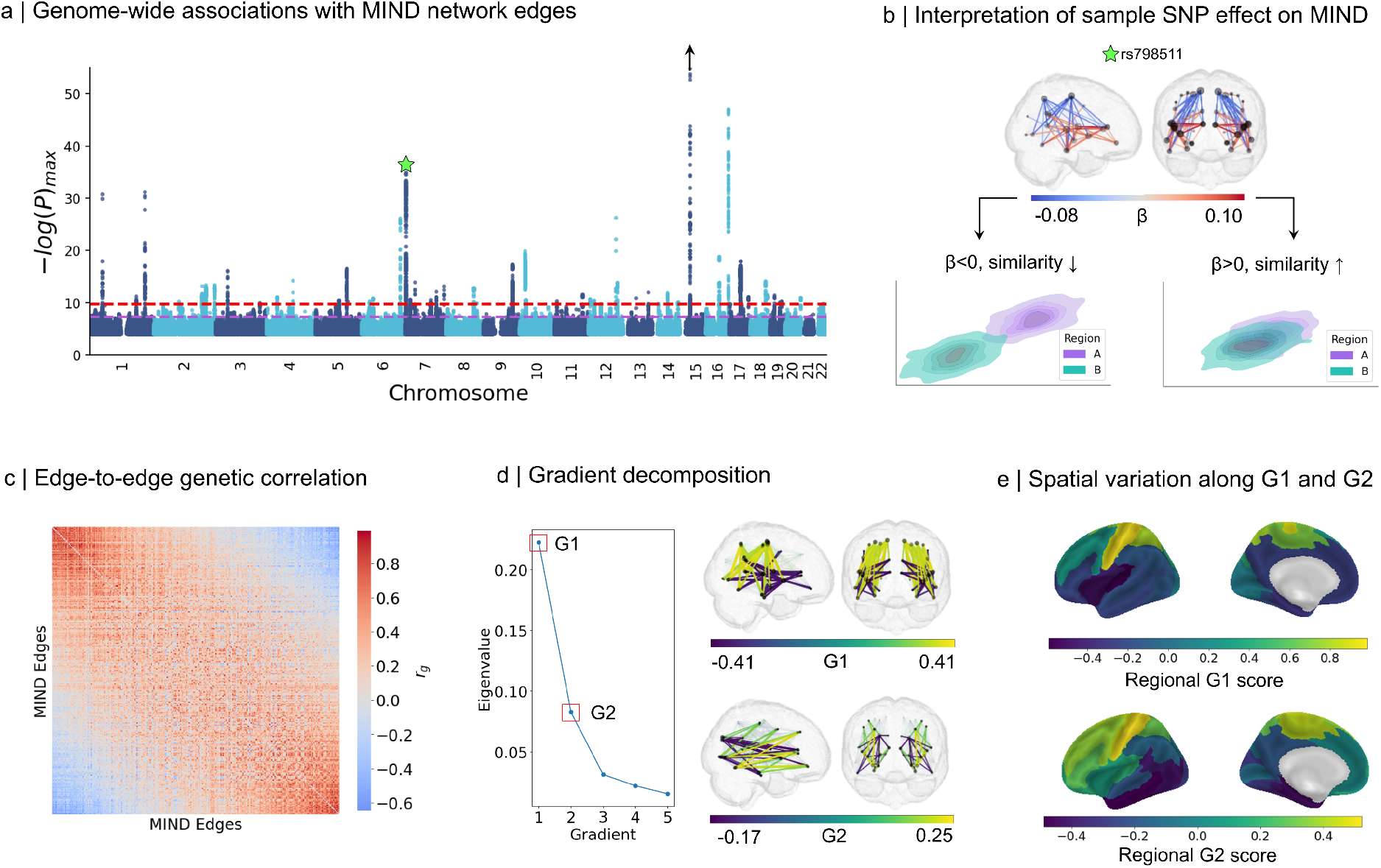
Edge-level genetic associations with structural similarity follow two underlying genetic gradients. A) Manhattan plot summarizing common variant associations with MIND networks across all 276 studied network edges. After clumping, 54 approximately-independent genomic loci were identified at experiment-wide significance (P*<*5e-08/276=1.8e-10) as being related to one or more MIND network edges. The *y*-axis indicates the most significant association of each variant across all edges. The purple line indicates genome-wide significance (P*<*5e-08), and the red line indicates experiment-wide significance (P*<*1.8e-10). The black arrow above one locus on chromosome 15 indicates that variants at that locus achieve higher significance than shown (up to *log*_10_(*P*) = 112), but are not shown here for ease of visualization. B) An illustration of the distributed network effects of a leading MIND-associated variant (indicated by a green star in panel A) and their interpretation in terms of a graphical representation of similarity. In the brain network plot, only edges achieving genome-wide significance are shown, and the size of each node is proportional to the number of associated edges reaching genome-wide significance. Coronal views show each edge repeated twice – once per intrahemispheric segment of each of the 23 regions (which span cortical areas on both hemispheres). C) Heatmap showing the genetic correlations between each MIND network edge, as measured by LD Score Regression (LDSC) (Bulik-Sullivan et al, 2015). The edges are ordered by the principal gradient of this edge-by-edge genetic similarity matrix (described in part D). D) The left plot shows the eigenvalues of the first five gradients of a decomposition of the edge-by-edge genetic correlation matrix shown in (C), with G1 and G2 highlighted. To the right, the top and bottom 5% of G1 and G2 are shown on the brain. As the decomposition was performed on an edge-by-edge matrix, each of the resulting gradients can be considered a network, with one value per edge. E) In order to interpret the spatial patterning of G1 and G2, the edge-level gradient values were mapped to the regional-level via gradient decomposition. For each gradient, the regional score indicates how similar each region is along the corresponding edge-level gradient; for example, regions with similar regional G1 scores share similar profiles of edge-level G1 values. Gradient-level regional scores are shown as heatmaps on the cortical surface.

MIND edge-associated genome loci exhibited both highly distributed effects on similarity across the network (e.g., *rs798511* shown in Fig. 2B) and highly specific effects on edges connected to a single cortical area (e.g., *rs492435*, shown in Extended Data Fig. 2A). However, on average, genetic variants that were more significantly associated with MIND at any edge were also associated with a higher number of edges (Extended Data Fig. 2B), suggesting a general tendency for distributed rather than localized effects of genetic variation on MIND network phenotypes.

### Gradients of genetically correlated inter-areal similarity

Given this evidence for distributed effects of localised genetic variation on similarity (or dis-similarity) of multiple pairs of cortical areas, we estimated the genetic correlations between all possible pairs of MIND edges, compiled as a {276 edge × 276 edge} genetic correlation matrix, *G* (Fig 2C). There were evidently many strong genetic correlations between edges in the network and we used gradient decomposition (Vos de Wael et al, 2020) to summarise genetic effects on MIND networks in terms of the first two gradients, G1 and G2 (Fig 2D), which together accounted for over 70% of the (co)variance in *G* (see Methods for details). As the gradient decomposition was performed on an edge-by-edge matrix, each of the resulting gradients can be considered a network, with each edge weighted by its coefficients on G1 or G2.

To further investigate the genetic architecture of these two principal gradients, we first estimated gradient-level phenotypes by the sum of MIND edges in each individual network weighted by their coefficients on each of G1 and G2, using methods previously described for analysis of whole brain gradients of gene expression (Dear et al, 2024). Then we performed two additional GWAS of the gradient-level phenotypes. We found that G1 and G2 were genetically independent of each other, evidenced by a non-significant genetic correlation between them (LDSC; *r*_*g*_=0.08, *P*=0.18). Both MIND gradients were more heritable than MIND edges (on average): 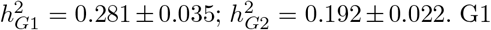 was significantly associated with 31 loci at the genome-wide threshold of *P* < 5 × 10^−8^; G2 was associated with a single locus at this threshold (see Extended Data Fig. 3A). This discrepancy between the number of associated loci for G1 and G2 was consistent with our finding that G1 was roughly ten times *less* polygenic than G2 (*π*_*G*1_=0.33%, *π*_*G*2_=3.7% using SBayesS; Extended Data Fig. 3C) (Zeng et al, 2018, 2021). This estimate of polygenicity for G1 was also substantially lower than the values reported for any of 206 measures of DTI-derived white-matter connectivity (0.8%≤ *π*_*DTI*_ ≤9.6%, median *π*=7.5%) from a recent GWAS (Wainberg et al, 2024), suggesting that it is influenced by an unusually concentrated set of genes compared to both G2 and other structural brain connectivity phenotypes. Additionally, we observed that G1 was under significantly stronger negative selection (*S*) compared to G2 (*S*_*G*1_=-0.58, *S*_*G*2_=-0.08 using SBayesS; Extended Data Fig. 3C). The combination of higher heritability (*h*^2^), lower polygenicity (*π*), and more negative *S* suggested that genetic influences on G1 were stronger, more genomically localized and evolutionarily conserved than genetic effects on G2.

### Gradients of genetically correlated cortical similarity are aligned with dual origins

The dual origin theory of cortex emerged from histological studies of phylogenetically primitive mammals and reptiles in the early 20th century (Abbie, 1940; Dart, 1934; Sanides, 1971; Pandya et al, 2015) which suggested that the complex 6-layer lamination of isocortex (a.k.a. neocortex) evolved as a function of distance from two conserved allocortical areas or primordial anlagen, the paleocortex and the archicortex, which have simpler 3-layer lamination. This theory has been revitalized to explain not only evolutionary trends in cytoarchitectonic differentiation but also the ontogeny of brain structure (García-Cabezas et al, 2019; Pandya et al, 2015), brain network architecture (Goulas et al, 2019), and the functionally specialised organization of dorsal and ventral sensory processing streams in the adult human cortex (Waymel et al, 2020; Meng et al, 2022). Dual origin theory is often assumed to have a neurogenetic basis, with recent research suggesting that spatial patterns of genetically-determined variation in human cortical structure are anatomically consistent with the cytoarchitectonic trends predicted from the pre-human evolution of cortex (Valk et al, 2020, 2022). However, the evidence is currently somewhat inconsistent and incomplete, and the genetic mechanisms underlying these expected trends have not been clearly articulated. For instance, the principal genetic gradient of regional cortical thickness did not align, as theoretically predicted, with maps of geodesic distance from either paleocortical or archicortical origins (Valk et al, 2020).

We tested the hypothesis that the two principal gradients of the MIND genetic correlation matrix, G1 and G2, were anatomically aligned with the two trends of cortical differentiation predicted by dual origin theory. To do this, we mapped the edge-level weights on each gradient to an areal patterning via a second round of gradient decomposition that mapped the two {276 edge × 276 edge} matrices of weights of each edge on G1 and G2 to two vectors representing the weight of each cortical area on each of the genetic gradients (Fig. 2E, Fig. 3). If dual origin theory is correct, we reasoned, areal weights on the two genetic gradients should be closely linked to geodesic distance from one of the two cortical origins, with more distant regions having more divergent G1 or G2 scores.

**Fig. 3.**
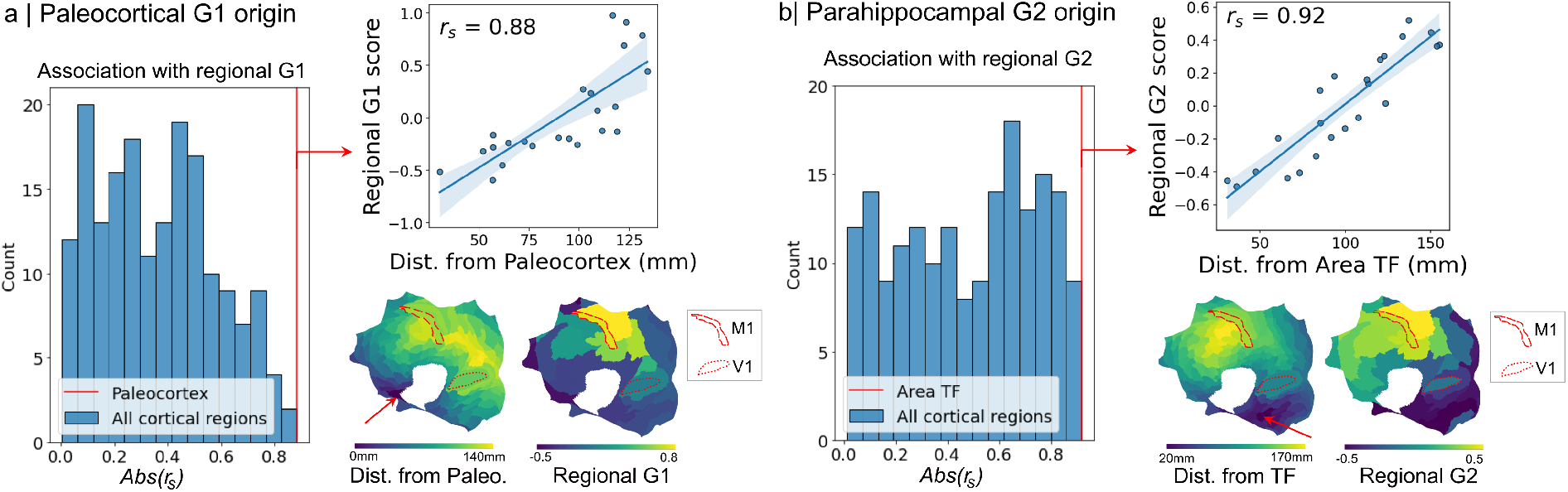
MIND genetics vary along the predicted trends of dual origin theory. A) The histogram indicates the alignment between the regional G1 scores and cortical maps of geodesic distance from each of 180 cortical regions from the HCP parcellation, plus separately-constructed maps of geodesic distance from the paleocortex and archicortex (the latter of which comprise multiple cortical regions) (*Abs*(*r*_*s*_) = Absolute Spearman’s correlation). The red line indicates the maximum association between regional G1 scores and region-specific distance, which corresponded to the paleocortical distance map. This association between regional G1 and paleocortical distance is shown to the right as a scatterplot. Beneath, the two cortical flatmaps show (left) the distance map from the paleocortex to each of 180 regions (bilaterally-averaged), with the paleocortex indicated by a red arrow, and (right) the regional G1 scores. B) The histogram is comparable to the one shown in part A, but for regional G2 scores. Distance from the parahippocampal area TF most closely aligned with regional G2 scores; this association between area TF and regional G2 is shown to the right as a scatterplot. The two cortical flatmaps show (left) the distance map from area TF to each of 180 regions (bilaterally-averaged), with area TF indicated by a red arrow.

The origin of the paleocortical trend is clearly identified as the piriform cortex, a small, discrete region adjacent to the anterior insula. As theoretically predicted, there was a strong correlation (Spearman’s *r*_*s*_=0.88) between a cortical map of G1 weights and a prior map of geodesic distance from the piriform cortex (Fig. 3A). To test the robustness and specificity of this association, we also calculated the correlations between areal G1 weights and maps of geodesic distance from each of the 180 (bilaterally-averaged) cortical areas designated by the HCP parcellation, which were coarse-grained to the 23 higher order areas for comparison. We found the cortical map of G1 scores was more strongly correlated with distance from the piriform cortex than with distance from any other cortical area (Fig. 3A), confirming that there was a uniquely strong alignment between the principal gradient of genetically determined (dis-)similarity between cortical areas and their distance from the paleocortical origin.

The origin of the archi-cortical trend in humans is more ambiguously defined, reflecting the more extensive, quasi-circular or limbic swathe of allocortex from the ventral hippocampus through parahippocampal and retrosplenial cortex to the dorsal remnants of hippocampus (indusium griseum and longitudinal striae) (Di Ieva et al, 2015). Therefore, to test the hypothesis that G2 was aligned with distance from the archi-cortical origin, we did not specify the origin *a priori* but calculated the correlations between areal G2 scores and geodesic distance from all 180 cortical regions in the HCP parcellation. We found that G2 scores were most closely aligned with distance from parahippocampal area TF (*r*_*s*_=0.92; Fig. 3B), an isocortical area adjacent to allocortical areas of the ventral hippocampus, which forms part of the evolutionary origin of the archi-cortical trend. However, areal G2 scores were not correlated with a prior map (Meng et al, 2022) of the archi-cortical trend in adult humans, which indexes distance from the entire limbic swathe of the archicortex (*r*_*s*_ = −0.03). In short, the second principal gradient of genetically coordinated patterning of cortical similarity, G2, was aligned with the archi-cortical trend (in contrast to the alignment of G1 with the paleocortical trend), broadly as predicted by dual origin theory. However, some ambiguities remain to be resolved, like the theoretically optimal origin for archi-cortical trend mapping in human MRI, before the putative identification of G2 with an archi-cortical trend of MIND similarity is as robust and specific as the identification of G1 MIND with the paleo-cortical trend of mammalian cortical evolution and expansion.

Dual origin theory was not discovered genetically, but by observing patterns of phenotypic variation of cortex across species. We therefore reasoned that a similar alignment to dual origin trends would be observed if MIND gradients were defined phenotypically, instead of genetically. To test this, we calculated the phenotypic covariance matrix *P*, representing the between-subject co-variation in MIND similarity for each possible pair of cortical areas, and found that this was strongly coupled with the genetic correlation matrix *G* (*r*_*s*_ = 0.85), as predicted by Cheverud’s conjecture (Cheverud, 1988) and previous work on structural MRI genetics (Stauffer et al, 2023) (Extended Data Fig. 4). The first two principal gradients derived from the phenotypic correlation matrix *P*, P1 and P2, were highly similar to G1 (*r*_*s*_ = 0.97) and G2 (*r*_*s*_ = 0.95), and their areal patterning was also correlated with areal G1 scores (*r*_*s*_ = 0.99) and G2 scores (*r*_*s*_ = 0.89), respectively.

As previously noted, G1 and G2 were genetically independent of each other (LDSC *r*_*g*_ = 0.08, *P* = 0.18). Spatially, however, the cortical patterning of areal weights on G1 and G2, i.e., the parcellated cortical maps of paleo-cortical and archi-cortical trends, were not fully orthogonal (*r*_*s*_ = 0.42, *P* = 0.05). Spatial co-location or convergence of these genetically independent cortical trends was more obvious for isocortical or koniocortical areas of higher laminar complexity at greater geodesic distance from the allocortical origins. For example, G1 and G2 originated from distinct allocortical locations (in piriform and parahippocampal cortex, respectively) but terminated in the same area of koniocortex (somatosensory cortex).

Overall these results support our prediction that the two principal gradients of genetically coordinated effects on MIND similarity networks should be anatomically aligned with the paleocortical (especially) and archicortical trends of dual origin theory (Goulas et al, 2019; Valk et al, 2020, 2022). As such, they show how dual origin theory can provide a framework for describing and genetically explaining two major trends of cortical differentiation in the human brain, in addition to its original focus on evolutionary progression of cortical structure across species.

### Genetic relationships between structural similarity and functional connectivity

These gradients of genetically coordinated structural similarity, as evident in human MRI data and predicted by dual origin theory, might be expected to have some consequences for functional organization of brain networks. MIND similarity can be compared directly to functional connectivity (FC) metrics, such as the correlation between resting-state fMRI time series, since both MIND and FC are relational measures that can be quantified between the same set of cortical areas. There is a large but inconclusive literature on the phenomenon of structure-function coupling of connectivity, typically estimated by the correlation of FC and structural connectivity metrics derived from axonal tractography of diffusion (d)MRI data, which is characterised by low subject-specificity and poor reliability across studies or processing measures (Zalesky et al, 2024; Zimmermann et al, 2018). Here, we tested the hypothesis that structural similarity (MIND) and functional connectivity (FC) metrics are robustly genetically coupled, and explored the extent to which structure-function coupling is differentially linked to the twin gradients of cortical genetic architecture.

To do this, we conducted an additional set of 276 GWAS on the same set of 276 edges as defined for the MIND networks, and in a subsample (*N*_*max*_=30,376) of the same UK Biobank participants; but now with each edge representing the correlation between regional mean fMRI time series recorded with the participant in a resting state (Fig. 4A). We also investigated the genetics of inter-individual variation in the aggregate strength of FC between areas weighted by the two MIND gradients G1 and G2. For each individual participant, we weighted and summed their FC matrix by the MIND-derived gradients G1 and G2, thereby projecting each FC network onto two gradients, FC1 and FC2, in the (directly-comparable) space of the paleo-cortical (G1) and archi-cortical (G2) trends in MIND similarity networks.

**Fig. 4.**
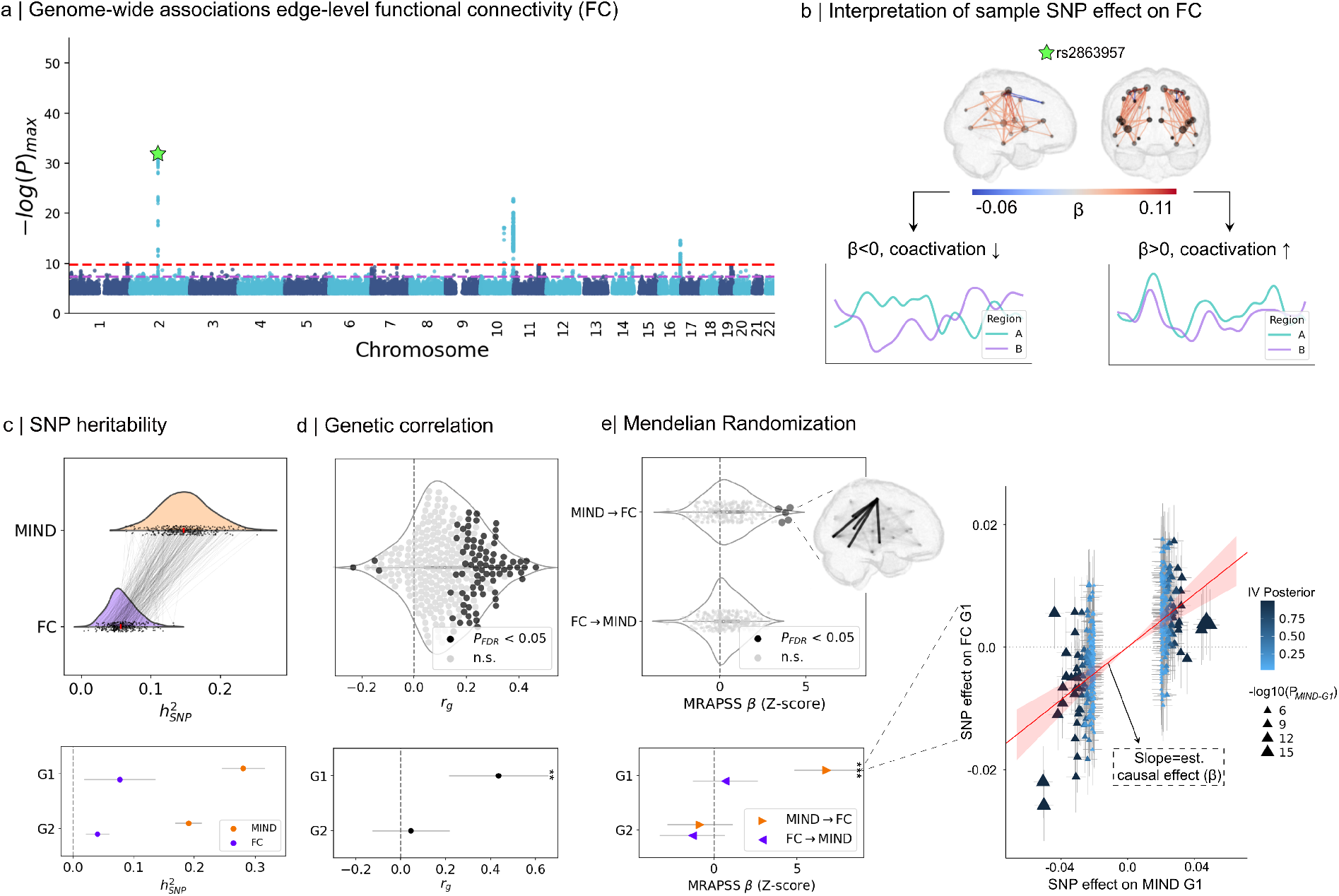
Genetic architecture of functional connectivity and its correlational and causal coupling with MIND similarity in paleocortical and archicortical trends. A) Manhattan plot summarizing common variant associations with functional connectivity (FC) network edges. After clumping, 4 approximately-independent genomic loci were associated at P < 1.8e-10 with one or more FC network edges. As in Fig. 2, the *y*-axis indicates the most significant association of each variant across all edges. The purple line indicates genome-wide significance (P < 5e-08), and the red line indicates experiment-wide significance (P < 1.8e-10). B) Illustration of the meaning of positive and negative GWAS *β* coefficients for a sample SNP in terms of functional connectivity. The brain network shows all edges positively or negatively associated to the SNP indicated by a green star in panel A; positive weights indicate greater positive FC (co-activation), and negative weights indicate reduced FC and co-activation, as shown schematically. C) SNP-based heritability estimates for MIND similarity and functional connectivity (FC). The top plot shows a histogram of heritability estimates at each of the 276 network edges estimated by high definition likelihood (HDL) (Ning et al, 2020). The bottom plot shows the mean SNP-based heritability (with 95% CI) for MIND and FC connectivity which have been weighted and summed along the MIND-derived G1 and G2 gradients. In other words, these are the heritability estimates of individual MIND or FC connectivity profiles projected into the comparable, lower-dimensional space of the MIND-derived gradients. D) Genetic correlations between MIND similarity and FC. The top plot shows the distribution of genetic correlations between MIND and FC over all (267) edges in the cortical network; solid points indicate 81 statistically significant genetic correlations between MIND and FC after FDR correction for *α* = 0.05. The bottom plot shows the gradient-level genetic correlations (and 95% CIs) between MIND and FC estimated from the GWAS of the first two gradients, G1 and G2. Genetic correlations were calculated using HDL (Ning et al, 2020) and replicated using LDSC (Bulik-Sullivan et al, 2015) (Fig. A1). E) Causal relationships between MIND similarity and functional connectivity. The top plot shows the distributions of *Z*-scored causal effects in both directions (MIND → FC and FC → MIND) for all (276) edges; solid points indicate genetic correlations that were statistically significant after FDR correction for *α* < 0.05. The inset brain network plot shows the location of all edges where MIND similarity had a significant causal effect on FC. The bottom plot shows estimated (forward and reverse) causal relationships between MIND and FC (with 95% CI) separately for both G1 and G2. The right plot shows the highly significant causal effect of structural similarity (MIND G1) on functional connectivity (FC1) in the paleocortical trend defined by the first gradient (G1) of the MIND genetic correlation matrix (Hu et al, 2022). This is illustrated by the significantly positive slope of the regression plot, which represents the amount of change expected for FC due to a corresponding change in MIND along G1.

At the phenotypic level of edges, MIND was more heritable than FC for 97.8% of edges, with an average SNP-based heritability nearly three times greater than that of FC (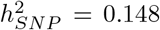 for MIND versus 0.058 for FC; see Fig. 4C). At the phenotypic level of gradients, FC1 was marginally (though not significantly) more heritable than FC2, (Fig. 4C; 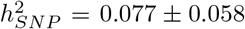 for G1, 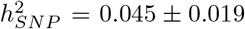 for G2); although both FC1 and FC2 had much lower heritabilities than G1 and G2, respectively (Fig. 4C). The relatively low heritability of functional connectivity phenotypes is in line with prior results (Roelfs et al, 2024) and demonstrates that FC is less strongly influenced by common genetic variation than MIND. In comparison to MIND, which showed substantial differences in both polygenicity (π) and negative selection (*S*) between G1 and G2, there were no significant differences between FC1 and FC2 on these parameters (*π*_*FC*1_ = 0.007 [0.004, 0.011], *π*_*FC*2_ = 0.01 [0.005, 0.017]; *S*_*FC*1_ = −0.52 [−0.96, −0.8], *S*_*FC*2_ = −1.09 [−1.30, −0.86]; Fig. A2).

Despite the relatively low heritability of functional connectivity phenotypes, we found solid evidence that they were genetically correlated with structural similarity phenotypes. Specifically, genetic correlations between MIND and FC edges were positive for 86% of network edges, with mean *r*_*g*_ = 0.12 (−0.23 ≤ *r*_*g*_ ≤ 0.46). There were significant positive genetic correlations at 28% of edges (*P*_*FDR*_ < 0.05), in contrast to significant negative correlations at about 1% of edges. Phenotypic correlations across individuals between MIND and FC edges were also generally positive (*r*_*s*_ *>* 0 for 83% of edges; *-*0.06 ≤ *r*_*s*_ ≤ *·* 0.13) and were moderately associated with the corresponding genetic correlations (*r*_*s*_ = 0.36,*P* < 0.001).

The genetic relationship between MIND and FC was clarified and strengthened through gradient-level comparison. On the paleocortical trend represented by G1, MIND G1 scores and FC1 scores were significantly genetically correlated at approximately the same level as the most strongly genetically correlated edges (HDL *r*_*g*_=0.44, *P* = 1.2e-04; LDSC *r*_*g*_ = 0.46, *P* = 6.5e-08). However, on the archicortical trend represented by G2, there was no significant genetic correlation between MIND G2 and FC2 scores (HDL *r*_*g*_ = 0.04, *P* = 0.61; LDSC *r*_*g*_ = 0.05, *P* = 0.76; Fig. 4D). This result suggests that the two genetically independent trends of cortical architecture, defined by structural MRI network analysis and aligned with dual origin theory based on cortical histology, have quite distinct relationships to brain functional activity.

We further investigated the potentially causal relationships between structural similarity and functional connectivity using Mendelian randomization (MR). We fit MR models for each MIND and FC network edge, as well as for the paired gradients G1/FC1 and G2/FC2, in both the forward (MIND ⟶ FC) and reverse (FC ⟶ MIND) directions using MR-APSS, a robust method that corrects for horizontal pleiotropy as well as the presence of overlapping samples (Hu et al, 2022, 2024).

At the level of edges, we found evidence for a significant forward causal relationship (MIND ⟶ FC) at multiple edges (*P*_*FDR*_ = 0.05 across 276 tests; Fig. 4E), but no evidence for significant reverse causal relationships. At the level of gradients, these findings came into sharper focus, showing a strong and specific forward causal relationship between MIND G1 and FC1 (Fig. 4E; *Z*-scored *β*=6.8, *P* = 1.4e-11); but no significant forward causal relationship between MIND G2 and FC2; and no significant reverse causal relationships between FC1 and G1 or FC2 and G2. The causal relationship of MIND G1 on FC1 significantly replicated when using MIND summary statistics from the (non-overlapping) replication cohort in a strict two-sample setting (IVW *β* = 0.19, *P* = 1.3e-05; Extended Data Fig. 5B), and was robust across a number of alternative MR methods and sensitivity analyses (Extended Data Fig. 5A, Fig. A3).

Taken together, these results suggest that the strong genetic correlation between functional connectivity and structural similarity in the paleocortical trend represented by G1 is the outcome of genetically driven effects on structural similarity causally constraining functional connectivity. This evidence aligns with prior work suggesting that brain morphology constrains brain function, but not vice versa (Pang et al, 2023; Valk et al, 2022). More generally, the concordant correlational and causal relationships between MIND and FC, at least in the paleo-cortical gradient, are consistent with a *homophilic* interpretation, whereby genetically-coordinated structural similarity is associated with (and/or causes) functional connectivity and is predictive of anatomical connectivity (Sebenius et al, 2024). Brain networks have previously been shown to demonstrate homophilic properties across diverse experimental scales and modalities (from neuronal cultures to human MRI scans) and across a wide range of similarity metrics (including topological assortativity, genomic co-expression, and cytoarchitectonic profile covariation) (Hansen et al, 2023; Bazinet et al, 2023; Akarca et al, 2021, 2022). Here, the alignment between MIND to FC provides genetic and causal support for these diverse observations and simulations of homophily as an important organizing principle of brain networks.

### Pleiotropic associations between dual origin gradients and biomedical traits

Structural and functional brain network organization is the outcome of neurodevelopmental processes and can be atypical in neurodevelopmental or neurodegenerative brain disorders (Menon, 2013; Vértes and Bullmore, 2015; Bullmore and Sporns, 2009). Recent GWAS results have provided some evidence for genetic correlations between brain network phenotypes and neuropsychiatric disorders, as well as pleiotropic associations between structural MRI phenotypes and a wide range of other biomedical traits (Grasby et al, 2020; Stauffer et al, 2023; Zhao et al, 2021; Alex et al, 2023; Kitzbichler et al, 2023; Sha et al, 2023). As such, we expected to observe pleiotropic associations between MIND gradients G1 and G2 (and their related functional connectivity gradients, FC1 and FC2) and a range of biomedical traits. In light of prior work theorizing that the paleocortical and archicortical trends of dual origin theory are differentially implicated in the pathogenesis of brain disorders (Giaccio, 2006), we anticipated that G1 and G2 might be genetically linked to distinct clinical diagnostic categories or health-related traits. We therefore estimated the genetic correlations between the dual origin gradients of structural similarity and functional connectivity and 6 neuropsychiatric disorders and 3 biomedical traits.

The paleocortical trend of structural similarity represented by G1 was globally genetically correlated with schizophrenia (SCZ; *r*_*g*_ = 0.08, *P* = 0.03 corrected for two gradient-level tests). Follow-up MR analysis provided some evidence for a causal effect of G1 on SCZ (MR-APSS *β*=0.037, *P* = 0.04 uncorrected); with no evidence for a causal effect in the reverse direction (Table A2). The archicortical trend represented by G2 was globally genetically correlated with multiple disorders or traits, including SCZ (*r*_*g*_ = −0.09, *P* = 0.02), major depressive disorder (MDD; *r*_*g*_ = −0.15, *P* = 6.3*e* − 05), attention deficit hyperactivity disorder (ADHD; *r*_*g*_ = −0.11, *P* = 0.02), C-reactive protein concentration in serum (CRP; *r*_*g*_ = 0.11, *P* = 0.004), and body mass index (BMI; *r*_*g*_ = −0.13, *P* = 1*e* − 04). Subsequent MR analysis demonstrated no forward causal effect of G2 on any disorders or traits (Table A2). However, there was strong evidence for a reverse causal effect of BMI on G2 (MR-APSS *β*=-0.15, *P* = 8.0*e* − 04 uncorrected) and suggestive evidence for a reverse causal effect of CRP on G2 (MR-APSS *β*=-0.07, *P* = 0.045 uncorrected). These results provide some preliminary support for the dual origin theory that the two trends of cortical differentiation are differentially implicated in pathogenesis of neuropsychiatric disorders.

It has been shown that global genetic correlations only incompletely capture the shared genetic effects on brain structure and risk of psychiatric disorders (Sha et al, 2023), due to the admixture of local correlations in discordant directions (both negative and positive), or noise introduced by large and unrelated portions of the genome. We therefore estimated a comprehensive set of local genetic correlations between Dual Origin gradients, clinical disorders, and biomedical traits (Fig. 5A-D) (Zhang et al, 2021). We observed a higher number of significant local genetic correlations as compared to global genetic correlations (86% of MIND-trait pairs had ≥1 significant local *r*_*g*_ versus 32% with significant global *r*_*g*_), with many local correlations in discordant directions (Fig. 5A). Although MIND gradient G1 had fewer significant global genetic correlations with diagnostic and biomedical traits than G2, it had more significant local genetic correlations. For example, by global genetic correlation analysis CRP, ADHD, and MDD were only significantly linked to G2, not G1; but all these traits had more significant local genetic correlations with G1 than with G2.

**Fig. 5.**
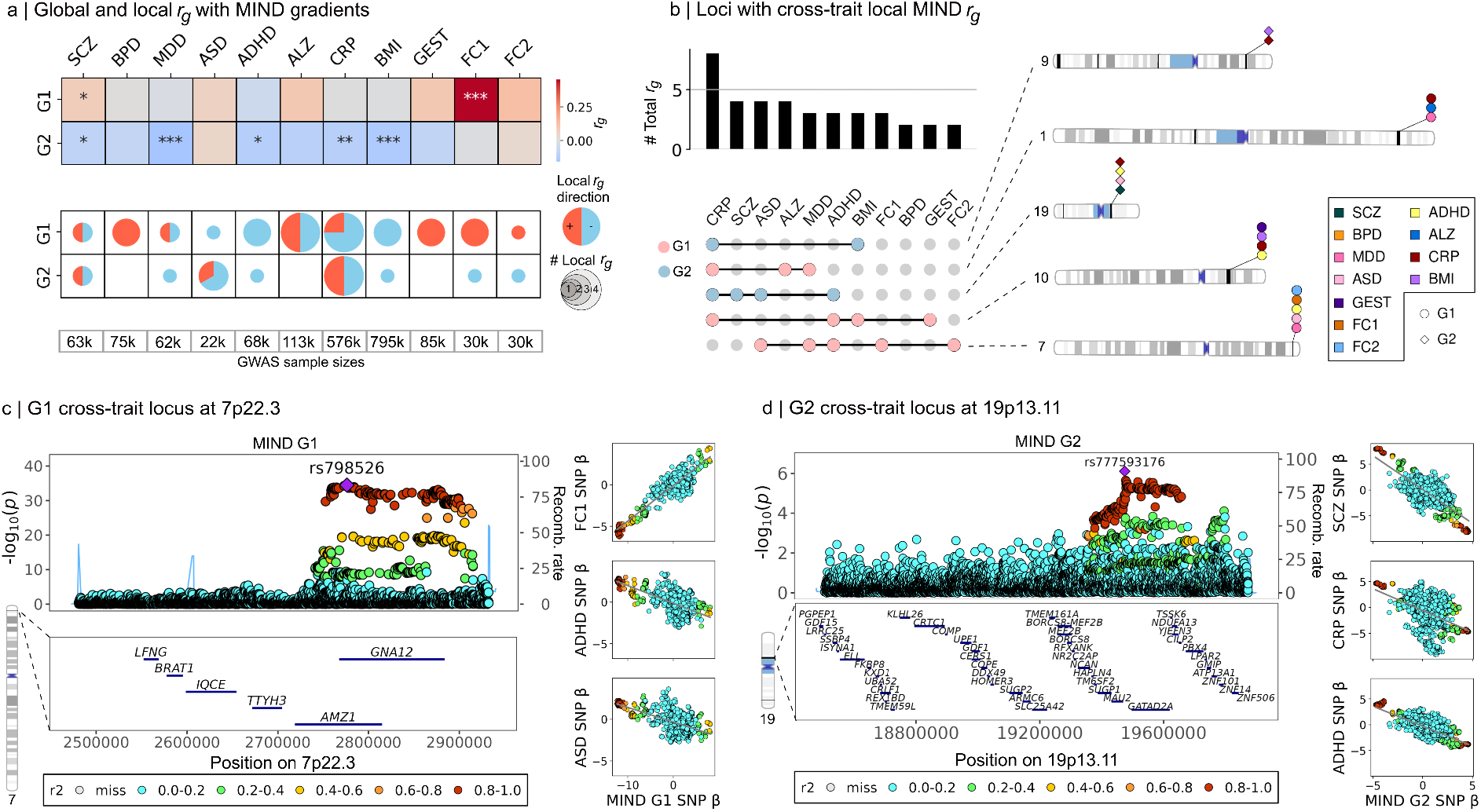
Pleiotropic associations between MIND genetic gradients, neuropsychiatric disorders and biomedical traits. A) Magnitude (and significance) of global genetic correlations (top row) and number and sign of significant local genetic correlations (middle row) between MIND genetic gradients, G1 and G2, and each of 6 neuropsychiatric disorders (SCZ, schizophrenia; BPD; bipolar disorder; MDD, major depressive disorder; ASD, autism spectrum disorder; ADHD, attention-deficit hyperactivity disorder; or ALZ, Alzheimer’s disease), each of 3 biomedical traits (CRP, C-reactive protein; BMI, body mass index; GEST, gestational age at birth), and both functional connectivity gradients, FC1 and FC2 (11 columns in total). The bottom row reports the size of all comparison GWAS, with estimated effective sample size reported for case-control traits and total sample size for quantitative traits (see Table A1). Global genetic correlations were FDR-corrected for the number of gradients (2 tests), and local genetic correlations were corrected for both the number of independent genomic regions and gradients (2, 262 × 2 = 4, 524 tests). \*\*\**P* < 0.001, **0.01 *<P* < 0.001, *0.01 *<P* < 0.05. B) Visualizations of the five cross-trait loci where ≥ 2 outcomes were significantly locally genetically associated with MIND G1 or G2. The barplot indicates the total number of significant local *r*_*g*_ s associated with each phenotype and either G1 or G2, of which the five cross-trait loci are highlighted below and in the chromosome maps to the right. C) Visualization of the shared local genetic effects between three comparison phenotypes (FC along G1, ADHD, and ASD) sharing a significant genetic correlation with MIND G1 at the locus spanning 2.5-2.9mb on chromosome 7p22.3. The locus plot shows the genes within the region and the variant-level association with MIND G1. The stacked scatter plots to the right illustrate the local genetic correlation by showing the (*Z*-scored) regression *β* values of the same regional variants for MIND and each of FC1, ADHD, and ASD. D) The same plots as in part (C), but for MIND G2 at a locus within 19p13.11 where it is locally genetically correlated with SCZ, CRP levels, ADHD, and (not shown) ASD.

Of the 24 genomic loci locally correlated with one or both of the dual origin gradients and one or more clinical diagnoses or biomedical traits, five loci were associated trans-diagnostically with multiple neuropsychiatric disorders (Fig. 5B). At one of these loci (within 7p22.3, shown in Fig. 5C), G1 was not only associated with multiple disorders (MDD, ADHD, and ASD) but was also associated strongly and positively with functional connectivity (FC1 local *r*_*g*_ ∼1, P < 1.0e-10; FC2 local *r*_*g*_ = 0.82, *P* = 3.2*e*−08). Colocalisation analysis (Dong et al, 2023) between G1 and FC1 suggested a high probability (0.928) of a shared causal variant, with the 95% credible set containing multiple eQTLs of *GNA12* and *AMZ1* such as *rs798526*, the SNP most strongly associated with G1, as highlighted in the gradient-level Manhattan plot in Extended Data Fig. 3A. Visual inspection showed that significant variants at this locus cleanly straddled the length of *GNA12* (Fig 5C). *GNA12*, encoding GCPR *α*-subunit 12, has been causally linked to several potentially relevant neurobiological processes including transmembrane signalling and regulating neuronal migration in the developing mouse cortex (Moers et al, 2008). This locus represents a specific and plausible set of genes pleiotropically associated with paleocortical trends in structural similarity and functional connectivity and trans-diagnostic risk of neuropsychiatric disorders.

Multiple pleiotropic associations were also observed for G2. For example, as shown in Fig. 5D, G2 was locally correlated with both a systemic inflammatory marker (CRP) and three neuropsychiatric disorders (ASD, ADHD, and SCZ) at one locus on 19p13.11, including *GATAD2*, which is a well-established schizophrenia risk gene linked to neurodevelopmental abnormalities (Merikangas et al, 2022; Werren et al, 2023). This locus represents a putatively distinct, neuroinflammatory mechanism for pleoiotropic association of archi-cortical structural similarity and trans-diagnostic risk of neurodevelopmental disorders.

## Conclusion

Here we investigated the genetic architecture of human cortical networks derived from structural MRI measures of the similarity between cortical areas. We concentrated our analysis of these data on predictions or questions arising from the dual origin theory of cortex, which originated in histological studies of non-human species published about 100 years ago but which had not yet been widely translated to (ante-mortem) genetic studies of human cortical structure, function, and disorder.

We found compelling evidence that the paleocortical trend predicted by dual origin theory conforms to the first principal axis of genetically coordinated variation in the human cortical architectome, gradient G1. There was also strong evidence that the second principal axis, gradient G2, was aligned with the theoretically predicted archi-cortical trend. These results suggest that the twin cortical trends of dual origin theory, originally defined to describe phylogenetic variation of the cortex across mammalian species, can also provide a scaffolding for analysis of genetically-mediated variation in structural MRI similarity networks measured within and between individual human brains. This new bridge between dual origin theory and human network neuroscience has allowed us to test some of the broader predictions of dual origin theory for human brain function and health. We have shown that these two genetically independent but anatomically converging gradients of cortical similarity are distinctly related to the genetic architecture of brain functional networks and pleiotropically associated with trans-diagnostic clusters of neuropsychiatric disorders and/or biomedical traits. These results generate many further questions about the detailed mechanisms of human cortical evolution and development of dual origin trends of inter-areal similarity, and their potential implications for a more neurogenetically grounded analysis of brain functional networks and pathogenic pathways to neuropsychiatric disorders. The GWAS summary statistics of MIND similarity network phenotypes (edges and gradients G1 and G2) and fMRI functional connectivity (edges and gradients FC1 and FC2) are made openly accessible to the research community to support further investigations into the origins and trends of the human cortical architectome.

## Methods

### Dataset

From the roughly 500,000 participants in the UK Biobank, our primary analysis was conducted using data from a total of 40,715 participants with magnetic resonance imaging (MRI) data available in 2023, of which 34,988 had the full set of MRI sequences and sufficient scan quality necessary to construct MIND networks. This study used the UKB genetic data that was previously quality-controlled and imputed before access as described by Bycroft et al (2018), and used largely the same set of participants and genetic inclusion criteria as described in Warrier et al (2023). Specifically, participants were included based on the following genetic criteria: 1) self-identification as ethnically white, 2) within ± 5 standard deviations of the mean along the first two genetic principal components, 3) matched genetic and reported sex, and 4) no excessive heterozygosity as reported by UKB. We included all SNPs with minor allele frequency greater than 0.1%, leading to the analysis of a maximum of 13,587,992 SNPs and 31,544 participants passing genetic inclusion criteria for the primary GWAS.

In 2024, the requisite MRI data for MIND network construction became available for an additional 21,447 UKB participants. This independent cohort formed our replication GWAS dataset. We applied the same genetic inclusion criteria to this validation cohort as for the original discovery cohort, leading to a maximum of 18,769 participants available for the replication GWAS.

### Magnetic Resonance Imaging (MRI) data

We used T1-weighted, diffusion-weighted, and resting-state functional MRI scans in the GWASs of UKB data. We additionally accessed T2-FLAIR images if available to facilitate surface reconstruction. The full imaging protocol can be found at: https://www.fmrib.ox.ac.uk/ukbiobank/protocol/V423092014.pdf and https://biobank.ctsu.ox.ac.uk/crystal/crystal/docs/brain_mri.pdf. In brief, the acquisition parameters for each type of data were the following, as described in Alfaro-Almagro et al (2018); Zhao et al (2022); Warrier et al (2023):

- **T1-weighted structural imaging (T1-w)**: 1.0 mm isotropic resolution, TR = 2,000 ms, TE = 2.01 ms, TI = 880 ms, flip angle = 8 degrees.
- **T2-weighted fluid-attenuated inversion recovery (T2-FLAIR)**: 1.0 × 1.0 × 1.1 mm resolution, TE = 395.0 ms, TR = 5,000 ms, TI = 1,800 ms.
- **Diffusion-weighted imaging (DWI)**: 2.0 mm isotropic resolution, MB = 3, R = 1, TE = 92 ms, TR = 3,600 ms, PF = 6/8, fat saturation, b = 0 s/mm**^2^** (5x + 3x phase-encoding reversed), b = 1,000 s/mm**^2^** (50x), b = 2,000 s/mm**^2^** (50x), 105 + 6 time-points (PA–AP).
- **Resting-state fMRI (rsfMRI)**: 2.4 mm isotropic resolution, flip angle = 52 degrees, fat saturation, TE = 39 ms, TR = 735 ms, scan duration = 6 min.

### Structural feature selection

Structural similarity necessarily depends on the measurements of brain structure on which it is based. Here, we estimated similarity based on a diverse set of interpretable macro- and micro-structural MRI features. We considered a total possible set of 13 structural MRI phenotypes which have previously been genetically analyzed in a recent GWAS study (Warrier et al, 2023): these included expansion-related phenotypes (e.g., surface area; SA, volume; Vol), cortical thickness (CT), curvature phenotypes (mean curvature; MC, Gaussian curvature; GC), standard diffusion-based phenotypes (mean diffusivity; MD, fractional anisotropy; FA) and neurite orientation dispersion and density imaging (NODDI) features from diffusion-weighted imaging (orientation dispersion index; ODI, isotropic volume fraction; ISOVF, and neurite density index; NDI, also known as intracellular-volume fraction; ICVF). From among these 13 phenotypes, we selected a final set of four MRI features as the basis for MIND similarity network estimation, each of which satisfied the following three criteria:

- **Local (vertex-level) interpretability**. MIND calculates structural similarity by comparing regional distributions of structural features measured at the level of individual vertices on the cortical surface. Therefore, we only considered features that were well-defined at the level of the individual vertex. For this reason, we excluded all expansion-based phenotypes including surface area and volume, which are only well-defined by aggregating (summing) information across vertices. Measured at the vertex level, these measurements are a direct function of the size and number of vertices used to reconstruct the surface mesh. Cortical thickness, curvature-based phenotypes, and diffusion-based phenotypes passed this criterion.
- **Genetic complementarity**. We aimed to integrate features capturing diverse genetic signals while avoiding the inclusion of multiple features that capture redundant signal. Warrier et al (2023) calculated the genetic correlations between each of the 13 globally-measured phenotypes and identified five genetically-defined clusters of structural features, representing: 1) expansion, 2) curvature, 3) thickness, 4) neurite density and orientation, and 5) water diffusion. To minimize redundancy while maximizing signal diversity, we included one feature from each cluster (except the expansion-related cluster excluded by the interpretability criterion).
- **SNP-based heritability**. CT was the only representative of the thickness cluster and had high heritability (*h*^2^ = 0.3). For each of the other three clusters, we selected a representative feature primarily by maximizing SNP-based heritability. For this reason, from the cluster of curvature phenotypes (MC and GC) we selected MC (*h*^2^=0.33 vs *h*^2^=0.19 for GC), and from the cluster of neurite density and orientation phenotypes (NDI, FA, and ODI), we selected NDI (*h*^2^=0.17 vs *h*^2^=0.11 for FA and *h*^2^=0.07 for ODI). The only exception to this preference for the highest heritability feature was for the water diffusion cluster (MD and ISOVF), where we selected MD (*h*^2^=0.15) over ISOVF (*h*^2^=0.24) as MD is more widely used in the neuroimaging community.

We settled on two macro-structural MRI phenotypes (mean curvature, MC; cortical thickness, CT) and two micro-structural phenotypes (mean diffusivity, MD; neurite density index, NDI). MC and CT were automatically defined at the vertex-level via default FreeSurfer outputs. MD and NDI were mapped from volumetric space (voxel resolution) to the vertex-level resolution of the cortical mesh via a surface-based registration in line with Human Connectome Project protocols, as described in Warrier et al (2023).

### MIND network construction

For each subject, the four MRI features were measured at every vertex on the subject-specific cortical surface mesh reconstruction, and aggregated over all vertices in each parcel of the widely-used 360-region HCP parcellation (Glasser et al, 2016), such that every region was defined by a four-dimensional distribution of the vertices contained within it. The similarity between each pair of these multivariate distributions was then estimated by morphometric inverse divergence (MIND) (Sebenius et al, 2023), which is the inverse of the Kullback-Leibler divergence between areal distributions, for all possible pairs of areas resulting in a single-subject {360 region x 360 region} structural similarity network. However, each of these networks contained 64,620 edges, which was too many to be tractable for edge-level GWAS analysis. We therefore coarse-grained the {360 region x 360 region} networks to {23 area x 23 area} networks comprising 276 edges for GWAS analysis, as illustrated in Figure 1. To perform this coarse-graining step, we leveraged the fact that the original HCP parcellation was derived in a hierarchical manner such that each of the 360 regions was assigned to one of 22 higher order cortical systems. Given the known cytoarchitectonic difference between primary motor cortex and primary somatosensory cortex (Geyer et al, 1997), we additionally split the higher-order sensorimotor system into its component motor and somatosensory cortical areas, leading to a total of 23 systems to which each of the 360 parcellated regions were assigned. Similarities within and between each of the 23 cortical systems were averaged for each participant, leading to the construction of subject-specific {23 area x 23 area} MIND networks.

### Functional connectivity estimation

Generating FC matrices from raw fMRI data involved three main steps: fMRI preprocessing, fMRI parcellation, and band-pass filtering. The first processing step was previously performed by the UKB and is described here (https://biobank.ctsu.ox.ac.uk/crystal/crystal/docs/brain_mri.pdf). Briefly, it includes motion correction using MCFLIRT (Jenkinson et al, 2002); grand-mean intensity normalisation of the entire 4D dataset by a single multiplicative factor; highpass temporal filtering (Gaussian-weighted least-squares straight line fitting, with sigma=50.0s); EPI unwarping; GDC unwarping; and structured artefact removal. The HCP parcellation was transformed from fsaverage standardised space to native space using surface-based non-linear registration. fMRI was linearly co-registered to the T1 image using FLIRT (Jenkinson et al, 2012). The resulting inverse transformation was used to map the T1-based parcellation into the fMRI space for the extraction of the average time series of each parcel. Finally, we focused on the physiologically relevant frequency range by using wavelet filtering that retained the BOLD oscillations in the frequency range 0.01–0.10 Hz (wavelet scales 2, 3 and 4).

### Genome-wide association analysis

For the primary GWAS on MIND network edges, in order to mitigate the effect of technical outliers, subjects were additionally excluded if they had values ± 5 standard deviations from the mean across any of the 276 MIND network edges, leading to a maximum of 30,524 participants. The same imaging inclusion criteria was applied to the replication cohort leading to a maximum of 18,769 subjects in the validation cohort.

For the subsequent gradient-level GWAS, subjects were also removed if they were ± 5 standard deviations from the mean subject-specific values of G1 and G2, leading to a maximum of 30,627 subjects in the discovery cohort. The same procedure for outlier control was applied to the corresponding GWAS of edges and gradients in the FC network, leading to a maximum of 30,376 and 29,715 participants, respectively, for these two phenotypic levels of GWAS.

In total, we conducted a total of 276 × 2 (MIND and FC edges) + 2 × 2 (MIND and FC gradients) = 556 genome-wide association studies (in the discovery cohort). In all GWAS, as in Warrier et al (2023), we standardized each phenotype to have a mean of zero and standard deviation of 1. We conducted all GWAS using FastGWA (Jiang et al, 2019) and as previously described in Warrier et al (2023) we incorporated the following covariates: age (years), age^2^, sex, age×sex, age^2^×sex, the first 40 genetic PCs as provided by the UKB, both mean and maximum framewise displacement (FD) corresponding to motion in the rs-fMRI scan, Euler Index, dummy variables encoding imaging center, and a dummy variable indicating whether T2 images were available for each subject, which may affect cortical surface reconstruction. After running edge-level GWAS, independent loci were identified via two-stage clumping using FUMA (Watanabe et al, 2017) with *P* < 0.05 at a threshold of *α* = 1.8*e* − 10 and default parameters of *r*^2^=0.6 and *r*^2^=0.1. The resulting loci were then merged if they fell within 250 kb of each other, thus reducing the number of loci from 69 to the final number of 54 reported in the main text. The same process was used to find independent loci for gradient-level GWAS, but with the a threshold of *α* = 5.8*e* − 08.

### Genetic correlations and causal analysis

To calculate the pairwise genetic correlations between MIND network edges, we used LD Score Regression (Bulik-Sullivan et al, 2015), with HapMap3 SNPs (The International HapMap 3 Consortium, 2010) and LD scores pre-computed from the 1KG European reference panel (The 1000 Genomes Project Consortium, 2015), due to its widespread use and fast runtime, which enabled 276*(276-1)=75,900 total genetic correlations needed to construct the pairwise genetic correlation matrix. Genetic correlations between MIND and FC were computed primarily using High-Definition Likelihood (HDL), and their precomputed reference panel of 1,029,876 quality-controlled UK Biobank imputed HapMap3 SNPs (Ning et al, 2020). This is more computationally expensive but has been shown to be a more powerful approach that yields comparatively more precise estimates of genetic correlations especially when SNP-heritabilities are low, as is the case with edge-level functional connectivity (Fig. 4C). Genetic correlations were compared with those derived from LD Score Regression and were highly consistent (Fig A1). SNP-based heritabilities for all MIND and FC phenotypes were additionally computed using HDL (Ning et al, 2020). Local genetic correlations between MIND and FC were computed using SUPERGNOVA (Zhang et al, 2021). Quasi-independent genomic regions were estimated and provided by the original SUPERGNOVA software. Colocalisation between MIND and FC was performed with coloc with default priors of *p*_1_ = 1*e*−04, *p*_2_ = 1*e* − 04, *p*_12_ = 1*e -* 05 (Giambartolomei et al, 2014; Wallace, 2020).

Mendelian randomization was conducted primarily using MR-APSS, a novel method which accounts for sample structure including sample overlap, and increases power through the inclusion of more genetic instruments without increased false positive rate (Hu et al, 2022). This method was particularly suitable for our analysis given a) sample overlap in the GWAS performed for FC and MIND, and b) few genetic instruments available at the standard significance threshold (*P* < 5*e* − 08) for many edges. Here we used an instrument-level cutoff of *P* < 5*e* − 04, which falls within the recommended range (Hu et al, 2022). In a recent benchmarking study of 16 MR methods, MR-APSS was the top-performing method in terms of statistical power, type 1 error, and robustness to genetic instrument selection (Hu et al, 2024) across all simulation scenarios. Nevertheless, we additionally performed several replication and sensitivity analyses of our main MIND-FC MR analysis including a) replicating the effect of G1 on FC1 using four standard two-sample MR methods (Inverse-Variance Weighted MR, Simple Median, Weighted Median, and MR-Egger) using MIND G1 (exposure) summary statistics derived from both the discovery and (fully non-overlapping) validation cohorts (Extended Data Fig. 5), b) testing robustness to MR-APSS model assumptions (Fig. A3B) and c) replicating the original MR-APSS results using a threshold of *P* < 5*e* − 05 (see Fig. A3A). Finally, we again used MR-APSS with an instrument-level cutoff of *P* < 5*e* − 04 for our analysis of causality between MIND G1 and G2 and the 9 biological traits analyzed for which we observed significant genetic correlations (results of sensitivity analyses of robustness to varying instrument thresholds are reported in Table A2).

To estimate the genetic parameters of polygenicity (*π*) and negative selection (*S*), we used SBayesS software from the GCTB toolkit (Zeng et al, 2018, 2021) with the same parameters as Wainberg et al (2024), including using the default ‘Sparse matrix (including MHC regions)’ as the LD matrix, and the flags `--chain-length 10000 --thin 10 --burn-in 2000`.

### Genetic gradient calculation

After the {276 edge x 276 edge} genetic correlation matrix *G* was compiled, we identified the first five genetic gradients using gradient decomposition as implemented by the BrainSpace package (Vos de Wael et al, 2020), using the code parameters `GradientMaps(kernel=‘normalized angle’, approach=‘dm’)`, and fit without any thresholding. The eigenvalues returned by diffusion mapping cannot be directly used to calculate the variance explained in the original data; using the approximation described by Dear et al (2024), the first two gradients (G1 and G2) explained 53.7% and 17.4% of variance in *G*. To ensure that these gradients were robust to the matrix decomposition methodology, we repeated the process using principal component analysis (PCA). Replicating the results from the gradient decomposition, the first two principal components explained 54.4% and 17.7% of the variance in the matrix, respectively, and were almost perfectly correlated with with G1 and G2 (*r*_*s*_ > 0.99 in both cases).

To map the network-level gradients to regional scores, we performed a second gradient decomposition of each of G1 and G2 using the same method. These regional scores were then compared to distance maps from the paleocortex, the archicortex, or any of the 180 (bilaterally-averaged) regions from the HCP parcellation. Each distance map was originally defined as the distance to each of the 180 bilaterally-averaged regions in the HCP parcellation, and was then coarse-grained via averaging within each of the 23 higher-order regions to facilitate a direct comparison to the regional gradient scores. For cortical areas from the HCP parcellation, distance matrices were calculated for left and right hemispheres independently, then averaged, using Connectome Workbench (Marcus et al, 2011). Distance maps from paleocortex and complete archicortex were downloaded from Meng et al (2022).

## Data availability

All summary statistics for the 276 MIND network edges and two MIND genetic gradients will be made publicly accessible on University of Cambridge servers upon publication. Summary statistics for schizophrenia (SCZ), major depressive disorder (MDD), attention-deficit hyperactivity disorder (ADHD), autism spectrum disorder (ASD), bipolar disorder (BPD), and Alzheimer’s disease (ALZ) can be downloaded from https://pgc.unc.edu/for-researchers/download-results/ (Trubetskoy et al, 2022; Wray et al, 2018; Demontis et al, 2023; Grove et al, 2019; Mullins et al, 2021; Wightman et al, 2021). Summary statistics for C-reactive protein concentration (CRP), Body Mass Index (BMI) and gestational duration (GEST) can be downloaded, respectively, from https://ftp.ebi.ac.uk/pub/databases/gwas/summarystatistics/GCST90029001-GCST90030000/GCST90029070/, https://portals.broadinstitute.org/collaboration/giant/index.php/GIANT consortium data files#2018 GIANT and UK BioBank Meta-analysis, and https://egg-consortium.org/gestational-duration-2019.html (Liu et al, 2019; Yengo et al, 2018; Locke et al, 2015; Said et al, 2022). Further information for each of the summary statistics is provided in Supplementary Table A1. Paleocortical and archicortical distance maps were downloaded from (Meng et al, 2022).

## Code and data

Code to calculate MIND networks can be accessed at https://doi.org/10.5281/zenodo.7974716. FastGWA was conducted with GCTA software v.1.94.1 (Yang et al, 2011). Colocalisation was performed using coloc v5.2.3 and R v4.3.1. `Genal` was used for for MR sensitivity analysis (Rivier et al, 2024). Both plink v1.9 and 2.0 were used in this study (Purcell et al, 2007). Gradient decomposition was performed using BrainSpace python package v0.1.10 (Vos de Wael et al, 2020). Data analysis and visualization was conducted using Python 3.6, 3.9, and 3.11. Locus plots were created using https://github.com/jrs95/geni.plots. UpSet plots were created using the code from Lex et al (2014). Phenogram chromosome maps were created using the software at https://visualization.ritchielab.org/phenograms/plot.

## Extended Data Figures

**Extended Data Figure 1.**
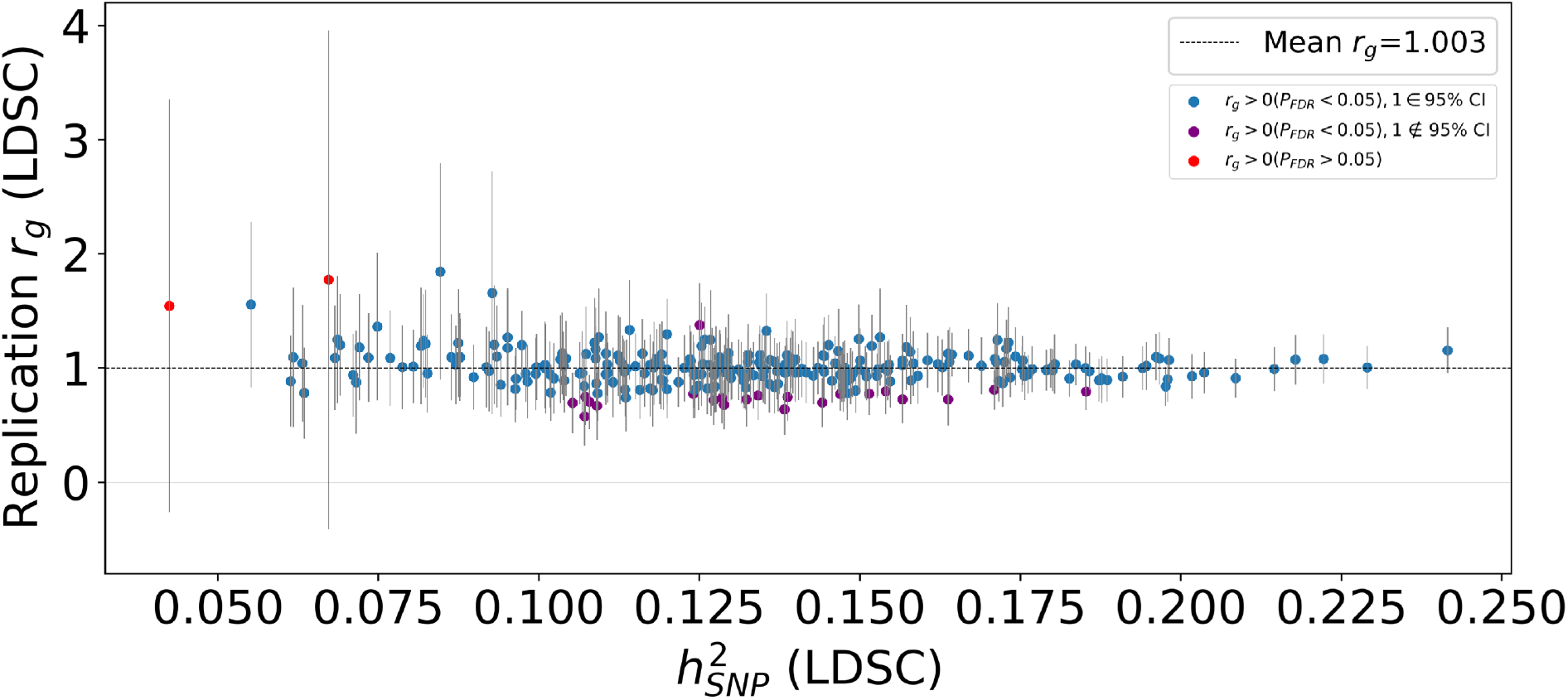
Replication of primary edge-level MIND GWAS. Scatterplot indicating the (LDSC) genetic correlation between the primary discovery GWAS, performed in *N* = 30, 524 subjects from the UK Biobank, and the replication GWAS from an additional *N* = 18, 769 from the UK biobank. The split between discovery and validation datasets naturally occurred due to data availability – data from the additional replication subjects became available in 2024, after the original study began in 2023. Each point in the scatterplot indicates one of the 276 total MIND network edges, with the *x*-axis representing its SNP-based heritability (LDSC-derived) and the *y*-axis showing the estimated genetic correlation between discovery and validation datasets, with 95% confidence intervals shown. The mean genetic correlation is shown as a dashed line (at *r*_*g*_ = 1.003). All but 274 of 276 edges had significantly positive genetic correlations across replication and discovery datasets, with two edges not reaching significance, demonstrating low heritability, shown in red. 254 of 276 edges (92%) had genetic correlations with confidence intervals overlapping with *r*_*g*_ = 1; the 22 edges where this was not the case are indicated in purple.

**Extended Data Figure 2.**
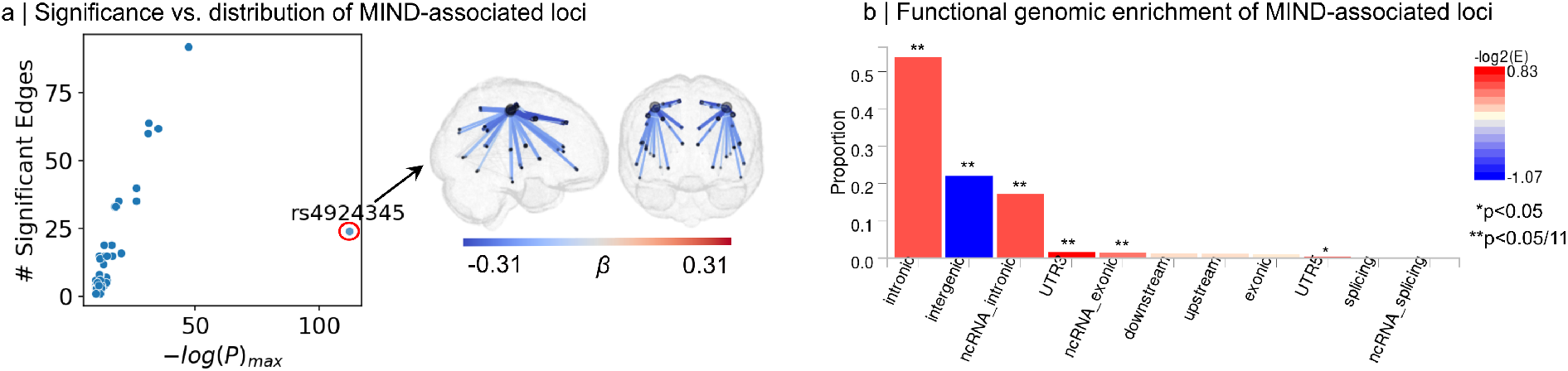
Distribution and enrichment of loci associated with MIND network edges. A) The left plot shows a scatterplot of the relationship between the maximum significance achieved by a given variant across all MIND network edges, and the number of edges at which it achieves significance. Each point represents the index variants for each of the 54 loci associated with at least one MIND network edge at an experiment-wide threshold of *P* < 1.8*e* 10. On average, the relationship between the maximum significance achieved and number of significance was highly positive, pointing to a strong general tendency for distributed network effects of MIND-associated loci. The one major outlier to this pattern was *rs4924345*; Occurring at chr15:39,639,898 (shown on the Manhattan plot in Fig. 2), this locus achieved the highest significance with MIND, but was associated with comparably few network edges. The correlation of maximum −*log*_10_(*P*) with the number of significantly-associated edges was *r*_*s*_ = 0.81. The brain plot on the right shows which edges were significantly associated with *rs4924345* at *P* < 5*e* − 08; as shown, they are exclusively focused on the somatosensory cortex. This result is consistent with previous findings showing associations with this variant and central sulcus morphology across multiple imaging-genetics datasets (van der Meer et al, 2021; Sun et al, 2022). B) Functional genomic enrichment plots computed by FUMA using ANNOVAR for ‘min-P’ values of genetic associations with MIND network edge connectivity across all 276 network edges. (Wang et al, 2010; Watanabe et al, 2017).

**Extended Data Figure 3.**
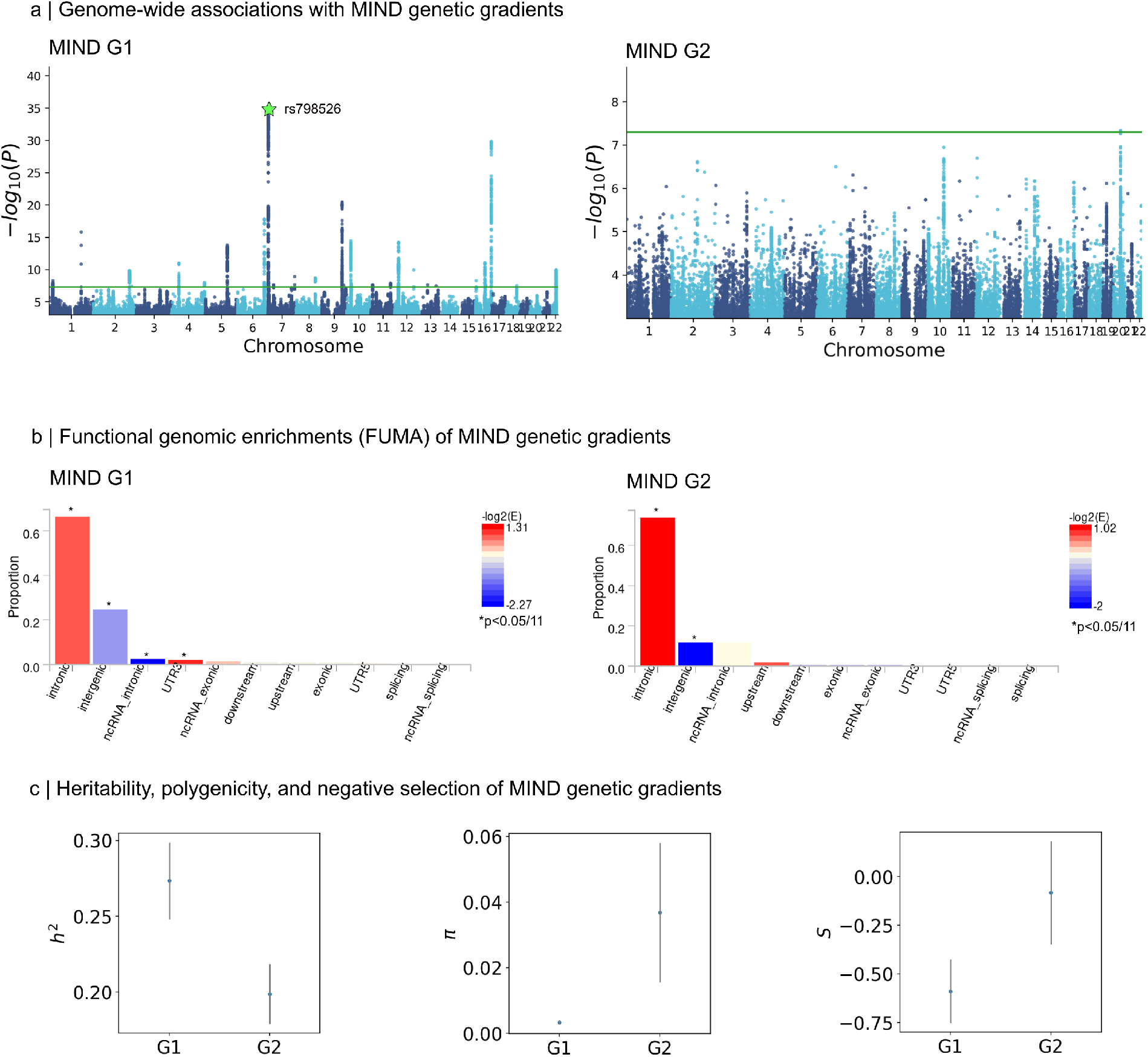
Gradient-level MIND genetic associations and enrichments. A) Manhattan plots showing the genetic associations with MIND, summarized via dot product with the edge-level weights of G1 and G2. The green line represents genome-wide significance of *P* < 5*e* − 08. In panel A, the green star highlights the top G1-associated variant, *rs798526*. B) Functional genomic enrichment plots computed by FUMA using ANNOVAR for summary statistics of MIND G1 and MIND G2 (Wang et al, 2010; Watanabe et al, 2017). C) Estimates of SNP-based heritability, polygenicity parameter *π*, and negative selection parameter *S* of the two principal MIND genetic gradients G1 and G2, calculated using SBayesS (Zeng et al, 2018). Shaded lines indicated 95% confidence intervals.

**Extended Data Figure 4.**
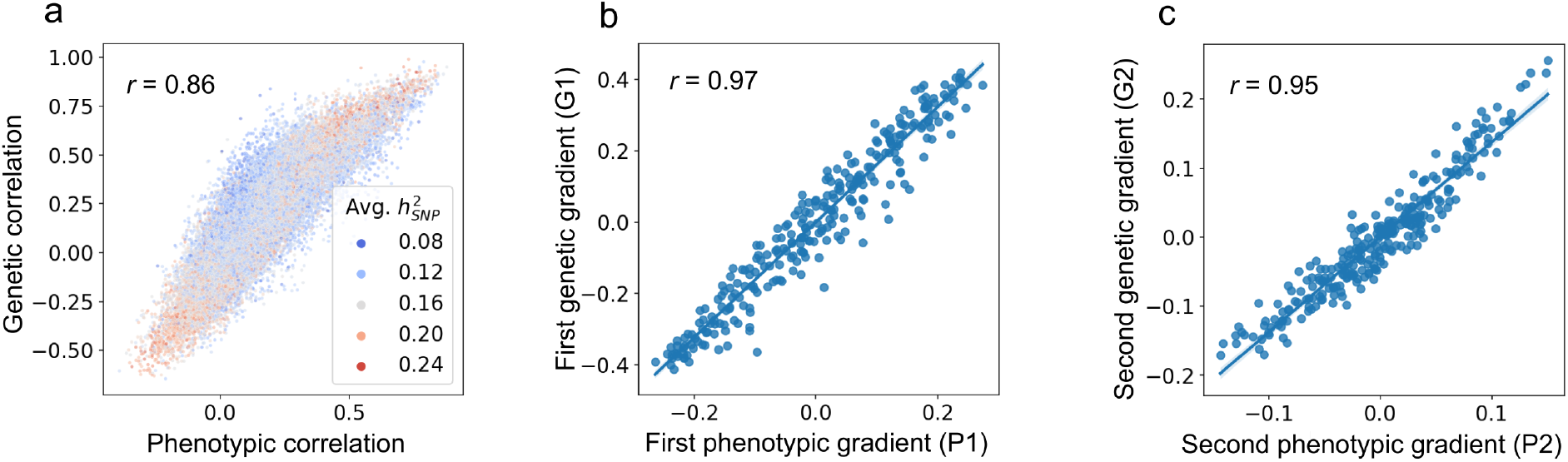
Genetic vs. phenotypic MIND network edge relationships. A) The association between phenotypic and genetic correlation between MIND network edges. Each point in the scatterplot represents a pair of edges. B-C) The first two phenotypic gradients (P1 and P2) defined by gradient decomposition of the edge-by-edge phenotypic correlation matrix (*P*) were highly similar to the first two genetic gradients (G1 and G2) derived from the edge-by-edge genetic correlation matrix (*G*).

**Extended Data Figure 5.**
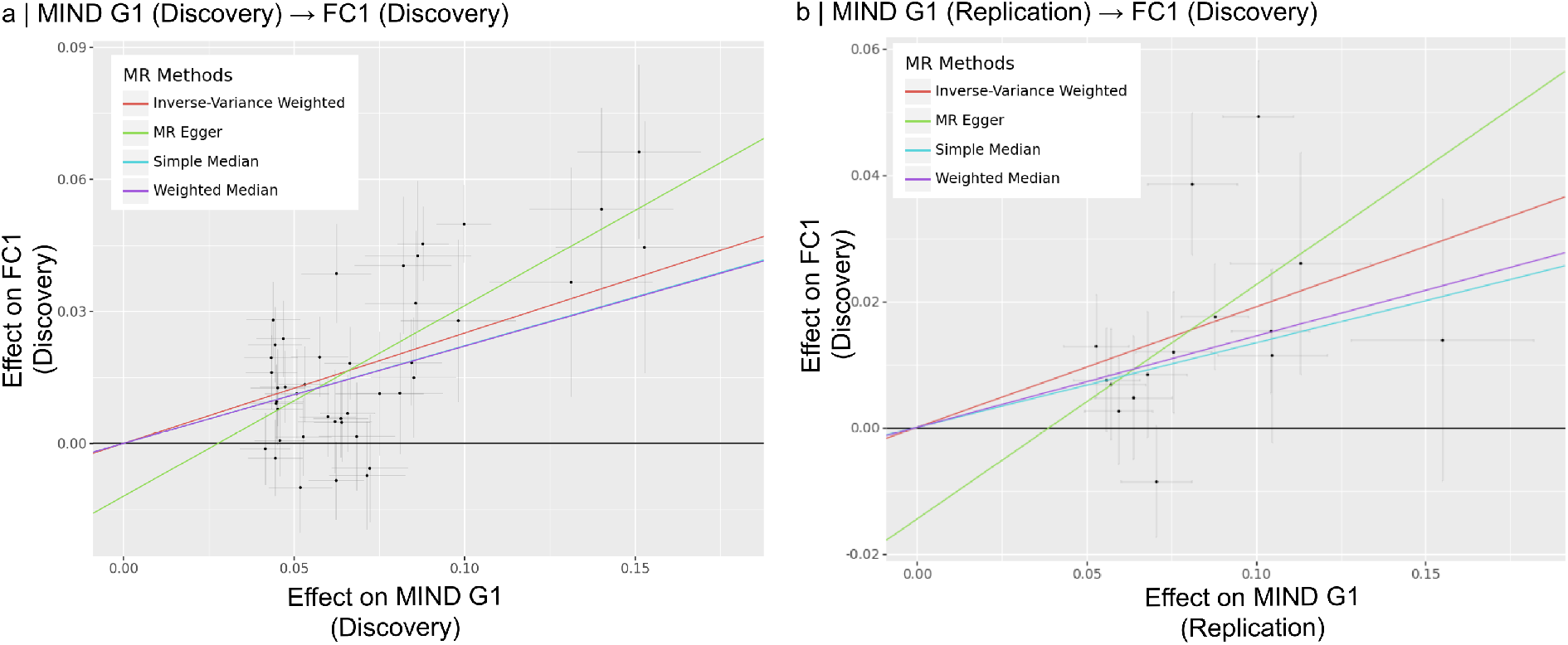
Replication of the causal effect of MIND on FC along G1. Results from four standard MR methods for estimating the causal effect of MIND on FC along G1, computed using MIND G1 summary statistics from the A) discovery cohort (*N*_*max*_ = 30, 627) and B) replication cohort (*N*_*max*_ = 18, 743). All results used the summary statistics for FC along G1 (FC1) from the discovery cohort. Thus, part B is a replication of these results using fully non-overlapping samples. Genetic instruments in both cases were selected using the standard strict threshold of *P* < 5*e* − 08. A) In this within-sample analysis, all four methods exhibited significant positive causal effects, with exact values as follows: IVW; *β* = 0.25,*P* = 7.5*e* − 15, MR Egger; *β* = 0.43,*P* = 1.3*e* − 04, Simple Median; *β* = 0.22,*P* = 8.2*e* − 08, Weighted Median; *β* = 0.22,*P* = 1.6*e* − 07. The Egger intercept was not significantly different from zero (-0.01, *P* = 0.07). B) Replicating these results in a strict two-sample framework with no subject overlap, all four methods again showed significant positive causal effects, with exact values as follows: IVW; *β* = 0.19,*P* = 1.3*e* − 05, MR Egger; *β* = 0.37,*P* = 4.4*e* − 02, Simple Median; *β* = 0.13,*P* = 7.9*e* − 03, Weighted Median; *β* = 0.14,*P* = 1.7*e* − 03. The Egger intercept was again not significantly different from zero (-0.01, *P* = 0.28).

## References

Abbie A (1940) Cortical lamination in the monotremata. Journal of Comparative Neurology 72:429–467. 10.1002/cne.900720302, URL 10.1002/cne.900720302

Akarca D, Vértes P, Bullmore E, et al (2021) A generative network model of neurodevelopmental diversity in structural brain organization. Nature Communications 12:4216. 10.1038/s41467-021-24430-z, URL 10.1038/s41467-021-24430-z

Akarca D, Dunn A, Hornauer P, et al (2022) Homophilic wiring principles underpin neuronal network topology in vitro. bioRxiv 10.1101/2022.03.09.483605, URL 10.1101/2022.03.09.483605, preprint

Alex AM, et al (2023) Genetic influences on the developing young brain and risk for neuropsychiatric disorders. Biological Psychiatry 93(10):905–920. 10.1016/j.biopsych.2023.01.013

Alfaro-Almagro F, et al (2018) Image processing and quality control for the first 10,000 brain imaging datasets from UK Biobank. NeuroImage 166:400–424. 10.1016/j.neuroimage.2017.10.012

Bassett DS, Sporns O (2017) Network neuroscience. Nature Neuroscience 20(3):353–364. 10.1038/nn.4502, URL 10.1038/nn.4502

Bazinet V, Hansen J, Vos de Wael R, et al (2023) Assortative mixing in micro-architecturally annotated brain connectomes. Nature Communications 14:2850. 10.1038/s41467-023-38585-4, URL 10.1038/s41467-023-38585-4

Beul SF, Grant S, Hilgetag CC (2015) A predictive model of the cat cortical connectome based on cytoarchitecture and distance. Brain Struct Funct 220(6):3167–3184. 10.1007/s00429-014-0849-y, URL 10.1007/s00429-014-0849-y

Beul SF, Barbas H, Hilgetag CC (2017) A predictive structural model of the primate connectome. Scientific reports 7(1):43176

Bulik-Sullivan B, Loh P, Finucane H, et al (2015) LD Score regression distinguishes confounding from polygenicity in genome-wide association studies. Nature Genetics 47:291–295. 10.1038/ng.3211

Bullmore E, Sporns O (2009) Complex brain networks: graph theoretical analysis of structural and functional systems. Nature Reviews Neuroscience 10:186–198. 10.1038/nrn2575, URL 10.1038/nrn2575

Bycroft C, et al (2018) The UK Biobank resource with deep phenotyping and genomic data. Nature 562:203–209. 10.1038/s41586-018-0579-z

Cheng R, Yin R, Zhao X, et al (2024) Genome-wide association study of brain functional and structural networks. Network Neuroscience 8(1):319–334. 10.1162/netn_a_00356, URL 10.1162/netn_a_00356

Cheverud JM (1988) A comparison of genetic and phenotypic correlations. Evolution 42(5):958–968. 10.1111/j.1558-5646.1988.tb02514.x

Chiang MC, Barysheva M, Shattuck DW, et al (2009) Genetics of brain fiber architecture and intellectual performance. Journal of Neuroscience 29(7):2212–2224. 10.1523/JNEUROSCI.4184-08.2009

Dart RA (1934) The dual structure of the neopallium: Its history and significance. Journal of Anatomy 69(Pt 1):3

Dear R, Wagstyl K, Seidlitz J, et al (2024) Cortical gene expression architecture links healthy neurodevelopment to the imaging, transcriptomics and genetics of autism and schizophrenia. Nature Neuroscience 27:1075–1086. 10.1038/s41593-024-01624-4, URL 10.1038/s41593-024-01624-4

Demontis D, Walters RK, Athanasiadis G, et al (2023) Genome-wide analyses of ADHD identify 27 risk loci, refine the genetic architecture and implicate several cognitive domains. Nature Genetics 55:198–208. 10.1038/s41588-022-01285-8, URL 10.1038/s41588-022-01285-8

Dennis EL, Thompson PM, Jahanshad N (2019) Chapter 8 - Genetics of brain networks and connectivity. In: Munsell BC, Wu G, Bonilha L, et al (eds) Connectomics. The Elsevier and MICCAI Society Book Series, Academic Press, p 155–179, 10.1016/B978-0-12-813838-0.00008-X, URL https://www.sciencedirect.com/science/article/pii/B978012813838000008X

Di Ieva A, Fathalla H, Cusimano MD, et al (2015) The indusium griseum and the longitudinal striae of the corpus callosum. Cortex 62:34–40. 10.1016/j.cortex.2014.06.016

Dong S, Zhao N, Spragins E, et al (2023) Annotating and prioritizing human non-coding variants with RegulomeDB v.2. Nature Genetics 55:724–726. 10.1038/s41588-023-01365-3

Elliott LT, Sharp K, Alfaro-Almagro F, et al (2018) Genome-wide association studies of brain imaging phenotypes in UK Biobank. Nature 562:210–216. 10.1038/s41586-018-0571-7, URL 10.1038/s41586-018-0571-7

Foo H, Thalamuthu A, Jiang J, et al (2021) Novel genetic variants associated with brain functional networks in 18,445 adults from the UK biobank. Scientific Reports 11(1):14633. 10.1038/s41598-021-94182-9, URL 10.1038/s41598-021-94182-9

Fu J, Zhang Q, Wang J, et al (2024) Cross-ancestry genome-wide association studies of brain imaging phenotypes. Nature Genetics 56:1110–1120. 10.1038/s41588-024-01766-y, URL 10.1038/s41588-024-01766-y

Gajwani M, Oldham S, Pang JC, et al (2023) Can hubs of the human connectome be identified consistently with diffusion MRI? Network Neuroscience 7(4):1326–1350. 10.1162/netn_a_00324

García-Cabezas M, Zikopoulos B, Barbas H (2019) The structural model: a theory linking connections, plasticity, pathology, development and evolution of the cerebral cortex. Brain Structure and Function 224(3):985–1008. 10.1007/s00429-019-01841-9, URL 10.1007/s00429-019-01841-9, epub 2019 Feb 9

Geyer S, Schleicher A, Zilles K (1997) The somatosensory cortex of human: cytoarchitecture and regional distributions of receptor-binding sites. Neuroimage 6(1):27–45

Giaccio RG (2006) The dual origin hypothesis: An evolutionary brain-behavior framework for analyzing psychiatric disorders. Neuroscience & Biobehavioral Reviews 30(4):526–550. 10.1016/j.neubiorev.2005.04.021, URL https://www.sciencedirect.com/science/article/pii/S0149763405001454

Giambartolomei C, et al (2014) Bayesian test for colocalisation between pairs of genetic association studies using summary statistics. PLOS Genetics 10.1371/journal.pgen.1004383

Glahn DC, Winkler AM, Kochunov P, et al (2010) Genetic control over the resting brain. Proceedings of the National Academy of Sciences 107(3):1223–1228. 10.1073/pnas.0909969107, URL https://www.pnas.org/doi/abs/10.1073/pnas.0909969107, https://www.pnas.org/doi/pdf/10.1073/pnas.0909969107

Glasser M, Coalson T, Robinson E, et al (2016) A multi-modal parcellation of human cerebral cortex. Nature 536:171–178. 10.1038/nature18933, URL 10.1038/nature18933

Goulas A, Margulies DS, Bezgin G, et al (2019) The architecture of mammalian cortical connectomes in light of the theory of the dual origin of the cerebral cortex. Cortex 118:244–261. 10.1016/j.cortex.2019.03.002, URL https://www.sciencedirect.com/science/article/pii/S0010945219301054, the Evolution of the Mind and the Brain

Grasby KL, et al (2020) The genetic architecture of the human cerebral cortex. Science 367:eaay6690. 10.1126/science.aay6690

Grove J, Ripke S, Als TD, et al (2019) Identification of common genetic risk variants for autism spectrum disorder. Nature Genetics 51:431–444. 10.1038/s41588-019-0344-8, URL 10.1038/s41588-019-0344-8

Hansen JY, Shafiei G, Voigt K, et al (2023) Integrating multimodal and multiscale connectivity blueprints of the human cerebral cortex in health and disease. PLOS Biology 21(9):e3002314. 10.1371/journal.pbio.3002314, URL 10.1371/journal.pbio.3002314

Hu X, Cai M, Xiao J, et al (2024) Benchmarking mendelian randomization methods for causal inference using genome-wide association study summary statistics. The American Journal of Human Genetics 10.1016/j.ajhg.2024.06.016, URL https://www.sciencedirect.com/science/article/pii/S0002929724002222

Hu X, et al (2022) Mendelian randomization for causal inference accounting for pleiotropy and sample structure using genome-wide summary statistics. Proceedings of the National Academy of Sciences 119(28):e2106858119. 10.1073/pnas.2106858119, URL 10.1073/pnas.2106858119

Jenkinson M, Bannister P, Brady M, et al (2002) Improved optimization for the robust and accurate linear registration and motion correction of brain images. Neuroimage 17(2):825–841. 10.1016/s1053-8119(02)91132-8

Jenkinson M, Beckmann CF, Behrens TEJ, et al (2012) Fsl. NeuroImage 62(2):782–790. 10.1016/j.neuroimage.2011.09.015

Jiang L, et al (2019) A resource-efficient tool for mixed model association analysis of large-scale data. Nature Genetics 51:1749–1755. 10.1038/s41588-019-0480-1

Kitzbichler MG, Martins D, Bethlehem RA, et al (2023) Two human brain systems micro-structurally associated with obesity. eLife 12:e85175. 10.7554/eLife.85175, URL 10.7554/eLife.85175

Lex A, Gehlenborg N, Strobelt H, et al (2014) UpSet: Visualization of Intersecting Sets. IEEE Transactions on Visualization and Computer Graphics 20(12):1983–1992. 10.1109/TVCG.2014.2346248, InfoVis ‘14

Liu X, Helenius D, Skotte L, et al (2019) Variants in the fetal genome near pro-inflammatory cytokine genes on 2q13 associate with gestational duration. Nature Communications 10:3927. 10.1038/s41467-019-11881-8, URL 10.1038/s41467-019-11881-8

Locke AE, Kahali B, Berndt SI, et al (2015) Genetic studies of body mass index yield new insights for obesity biology. Nature 518:197–206. 10.1038/nature14177, URL 10.1038/nature14177

Luppi A, Gellersen H, Liu Z, et al (2024) Systematic evaluation of fMRI data-processing pipelines for consistent functional connectomics. Nature Communications 15:4745. 10.1038/s41467-024-48781-5, URL 10.1038/s41467-024-48781-5

Marcus D, Harwell J, Olsen T, et al (2011) Informatics and data mining: Tools and strategies for the human connectome project. Frontiers in Neuroinformatics 5:4. 10.3389/fninf.2011.00004

van der Meer D, et al (2021) The genetic architecture of human cortical folding. Science Advances 7:eabj9446. 10.1126/sciadv.abj9446, URL 10.1126/sciadv.abj9446

Meng Y, Yang S, Xiao J, et al (2022) Cortical gradient of a human functional similarity network captured by the geometry of cytoarchitectonic organization. Communications Biology 5:1152. 10.1038/s42003-022-04148-4, URL 10.1038/s42003-022-04148-4

Menon V (2013) Developmental pathways to functional brain networks: emerging principles. Trends in Cognitive Sciences 17(12):627–640. 10.1016/j.tics.2013.09.015

Merikangas AK, Shelly M, Knighton A, et al (2022) What genes are differentially expressed in individuals with schizophrenia? A systematic review. Molecular Psychiatry 27:1373–1383. 10.1038/s41380-021-01420-7, URL 10.1038/s41380-021-01420-7

Moers A, Nürnberg A, Goebbels S, et al (2008) Galpha12/Galpha13 deficiency causes localized overmigration of neurons in the developing cerebral and cerebellar cortices. Molecular and Cellular Biology 28(5):1480–1488. 10.1128/MCB.00651-07, URL 10.1128/MCB.00651-07, epub 2007 Dec 17

Mullins N, Forstner AJ, O’Connell KS, et al (2021) Genome-wide association study of more than 40,000 bipolar disorder cases provides new insights into the underlying biology. Nature Genetics 53:817–829. 10.1038/s41588-021-00857-4, URL 10.1038/s41588-021-00857-4

Ning Z, Pawitan Y, Shen X (2020) High-definition likelihood inference of genetic correlations across human complex traits. Nature Genetics 52:859–864. 10.1038/s41588-020-0653-y, URL 10.1038/s41588-020-0653-y

Pandya D, Petrides M, Cipolloni PB (2015) Cerebral Cortex: Architecture, Connections, and the Dual Origin Concept. Oxford University Press

Pang J, Aquino K, Oldehinkel M, et al (2023) Geometric constraints on human brain function. Nature 618:566–574. 10.1038/s41586-023-06098-1, URL 10.1038/s41586-023-06098-1

Paquola C, Vos de Wael R, Wagstyl K, et al (2019) Microstructural and functional gradients are increasingly dissociated in transmodal cortices. PLOS Biology 17(5):e3000284. 10.1371/journal.pbio.3000284, URL 10.1371/journal.pbio.3000284

Purcell S, Neale B, Todd-Brown K, et al (2007) Plink: a tool set for whole-genome association and population-based linkage analyses. American Journal of Human Genetics 81(3):559–575. 10.1086/519795

Rivier CA, Clocchiatti-Tuozzo S, Huo S, et al (2024) Genal: A Python Toolkit for Genetic Risk Scoring and Mendelian Randomization. medRxiv 10.1101/2024.05.23.24307776, URL 10.1101/2024.05.23.24307776

Roelfs D, van der Meer D, Alnæs D, et al (2024) Genetic overlap between multivariate measures of human functional brain connectivity and psychiatric disorders. Nature Mental Health 2:189–199. 10.1038/s44220-023-00190-1, URL 10.1038/s44220-023-00190-1

Said S, Pazoki R, Karhunen V, et al (2022) Genetic analysis of over half a million people characterises Creactive protein loci. Nature Communications 13:2198. 10.1038/s41467-022-29650-5, URL 10.1038/s41467-022-29650-5

Sanides F (1971) Functional architecture of motor and sensory cortices in primates in the light of a new concept of neocortex development. In: Advances in Primatology. Academic Press, p 137–208

Schutte NM, Hansell NK, de Geus EJC, et al (2013) Heritability of resting state EEG functional connectivity patterns. Twin Research and Human Genetics 16(5):962–969. 10.1017/thg.2013.55

Sebenius I, Seidlitz J, Warrier V, et al (2023) Robust estimation of cortical similarity networks from brain MRI. Nature Neuroscience 26(8):1461–1471. 10.1038/s41593-023-01376-7

Sebenius I, Dorfschmidt L, Seidlitz J, et al (2024) Structural mri of brain similarity networks. Nature Reviews Neuroscience 10.1038/s41583-024-00882-2

Seidlitz J, Váša F, Shinn M, et al (2018) Morphometric similarity networks detect microscale cortical organization and predict inter-individual cognitive variation. Neuron 97(1):231–247.e7. 10.1016/j.neuron.2017.11.039, URL 10.1016/j.neuron.2017.11.039

Sha Z, Warrier V, Bethlehem RA, et al (2023) The overlapping genetic architecture of psychiatric disorders and cortical brain structure. bioRxiv 10.1101/2023.10.05.561040, URL 10.1101/2023.10.05.561040

Smith SM, Douaud G, Chen W, et al (2021) An expanded set of genome-wide association studies of brain imaging phenotypes in UK biobank. Nature Neuroscience 24:737–745. 10.1038/s41593-021-00826-4, URL 10.1038/s41593-021-00826-4

Stauffer EM, Bethlehem RAI, Dorfschmidt L, et al (2023) The genetic relationships between brain structure and schizophrenia. Nature Communications 14:7820. 10.1038/s41467-023-43567-7

Sun BB, Loomis SJ, Pizzagalli F, et al (2022) Genetic map of regional sulcal morphology in the human brain from UK biobank data. Nature Communications 13:6071. 10.1038/s41467-022-33829-1, URL 10.1038/s41467-022-33829-1

The 1000 Genomes Project Consortium (2015) A global reference for human genetic variation. Nature 526:68–74. 10.1038/nature15393

The International HapMap 3 Consortium (2010) Integrating common and rare genetic variation in diverse human populations. Nature 467:52–58. 10.1038/nature09298

Tissink E, Werme J, de Lange SC, et al (2023) The genetic architectures of functional and structural connectivity properties within cerebral resting-state networks. eNeuro 10(4):ENEURO.0242–22.2023. 10.1523/ENEURO.0242-22.2023, URL 10.1523/ENEURO.0242-22.2023

Tissink EP, Shadrin AA, van der Meer D, et al (2024) Abundant pleiotropy across neuroimaging modalities identified through a multivariate genome-wide association study. Nature Communications 15:2655. 10.1038/s41467-024-46817-4, URL 10.1038/s41467-024-46817-4

Trubetskoy V, Pardiñas AF, Qi T, et al (2022) Mapping genomic loci implicates genes and synaptic biology in schizophrenia. Nature 604:502–508. 10.1038/s41586-022-04434-5, URL 10.1038/s41586-022-04434-5

Valk S, Xu T, Paquola C, et al (2022) Genetic and phylogenetic uncoupling of structure and function in human transmodal cortex. Nature Communications 13:2341. 10.1038/s41467-022-29886-1, URL 10.1038/s41467-022-29886-1

Valk SL, et al (2020) Shaping brain structure: Genetic and phylogenetic axes of macroscale organization of cortical thickness. Science Advances 6:eabb3417. 10.1126/sciadv.abb3417

Vértes PE, Bullmore ET (2015) Annual research review: Growth connectomics–the organization and reorganization of brain networks during normal and abnormal development. Journal of Child Psychology and Psychiatry 56(3):299–320. 10.1111/jcpp.12365, URL 10.1111/jcpp.12365

Vos de Wael R, Benkarim O, Paquola C, et al (2020) BrainSpace: a toolbox for the analysis of macroscale gradients in neuroimaging and connectomics datasets. Communications Biology 3:103. 10.1038/s42003-020-0794-7, URL 10.1038/s42003-020-0794-7

Wainberg M, Forde NJ, Mansour S, et al (2024) Genetic architecture of the structural connectome. Nature Communications 15(1):1962. 10.1038/s41467-024-46023-2, URL 10.1038/s41467-024-46023-2

Wallace C (2020) Eliciting priors and relaxing the single causal variant assumption in colocalisation analyses. PLOS Genetics 10.1371/journal.pgen.1008720

Wang J, He Y (2024) Toward individualized connectomes of brain morphology. Trends in Neurosciences 47(2):106–119. 10.1016/j.tins.2023.11.011, URL https://www.sciencedirect.com/science/article/pii/S0166223623002734

Wang K, Li M, Hakonarson H (2010) ANNOVAR: functional annotation of genetic variants from high-throughput sequencing data. Nucleic Acids Research 38(16):e164. 10.1093/nar/gkq603, URL 10.1093/nar/gkq603

Warrier V, Stauffer EM, Huang QQ, et al (2023) Genetic insights into human cortical organization and development through genome-wide analyses of 2,347 neuroimaging phenotypes. Nature Genetics 55:1483–1493. 10.1038/s41588-023-01475-y, URL 10.1038/s41588-023-01475-y

Watanabe K, Taskesen E, van Bochoven A, et al (2017) Functional mapping and annotation of genetic associations with FUMA. Nature Communications 8:1826. 10.1038/s41467-017-01261-5, URL 10.1038/s41467-017-01261-5

Waymel A, Friedrich P, Bastian PA, et al (2020) Anchoring the human olfactory system within a functional gradient. NeuroImage 216:116863. 10.1016/j.neuroimage.2020.116863, URL https://www.sciencedirect.com/science/article/pii/S1053811920303499

Wei Y, Scholtens LH, Turk E, et al (2018) Multiscale examination of cytoarchitectonic similarity and human brain connectivity. Network Neuroscience 3(1):124–137. 10.1162/netn_a_00057

Werren EA, Guxholli A, Jones N, et al (2023) De novo variants in GATAD2A in individuals with a neurodevelopmental disorder: GATAD2A-related neurodevelopmental disorder. HGG Advances 4(3):100198. 10.1016/j.xhgg.2023.100198, URL 10.1016/j.xhgg.2023.100198

Wightman DP, Jansen IE, Savage JE, et al (2021) A genome-wide association study with 1,126,563 individuals identifies new risk loci for Alzheimer’s disease. Nature Genetics 53:1276–1282. 10.1038/s41588-021-00921-z, URL 10.1038/s41588-021-00921-z

Winkler T, Day F, Croteau-Chonka D, et al (2014) Quality control and conduct of genome-wide association meta-analyses. Nature Protocols 9(6):1192–1212. 10.1038/nprot.2014.071, URL 10.1038/nprot.2014.071

Wray NR, Ripke S, Mattheisen M, et al (2018) Genome-wide association analyses identify 44 risk variants and refine the genetic architecture of major depression. Nature Genetics 50:668–681. 10.1038/s41588-018-0090-3, URL 10.1038/s41588-018-0090-3

Yang J, Lee SH, Goddard ME, et al (2011) GCTA: a tool for genome-wide complex trait analysis. American Journal of Human Genetics 88(1):76–82. 10.1016/j.ajhg.2010.11.011, URL 10.1016/j.ajhg.2010.11.011

Yengo L, Sidorenko J, Kemper KE, et al (2018) Meta-analysis of genome-wide association studies for height and body mass index in 700,000 individuals of European ancestry. Human Molecular Genetics 27(20):3641–3649. 10.1093/hmg/ddy271, URL 10.1093/hmg/ddy271

Zalesky A, Sarwar T, Tian Y, et al (2024) Predicting an individual’s functional connectivity from their structural connectome: Evaluation of evidence, recommendations, and future prospects. Network Neuroscience 10.1162/netn_a_00400, URL 10.1162/netn_a_00400

Zeng J, de Vlaming R, Wu Y, et al (2018) Signatures of negative selection in the genetic architecture of human complex traits. Nature Genetics 50(5):746–753. 10.1038/s41588-018-0101-4

Zeng J, Xue A, Jiang L, et al (2021) Widespread signatures of natural selection across human complex traits and functional genomic categories. Nature Communications 12:1164. 10.1038/s41467-021-21446-3

Zhang Y, Lu Q, Ye Y, et al (2021) SUPERGNOVA: local genetic correlation analysis reveals heterogeneous etiologic sharing of complex traits. Genome Biology 22:262. 10.1186/s13059-021-02478-w, URL 10.1186/s13059-021-02478-w

Zhao B, Ibrahim JG, Li Y, et al (2019) Heritability of regional brain volumes in large-scale neuroimaging and genetic studies. Cerebral Cortex 29(7):2904–2914. 10.1093/cercor/bhy157

Zhao B, Li T, Smith SM, et al (2022) Common variants contribute to intrinsic human brain functional networks. Nature Genetics 54:508–517. 10.1038/s41588-022-01039-6, URL 10.1038/s41588-022-01039-6

Zhao B, et al (2021) Common genetic variation influencing human white matter microstructure. Science 372(6547):eabf3736. 10.1126/science.abf3736, URL 10.1126/science.abf3736

Zimmermann J, Griffiths J, Schirner M, et al (2018) Subject specificity of the correlation between large-scale structural and functional connectivity. Network Neuroscience 3(1):90–106. 10.1162/netn_a_00055, URL 10.1162/netn_a_00055

