## Supplementary Figures A1-A3, Supplementary Tables A1-A2 for "The genetic architecture of cortical similarity networks"

### Supplementary Information

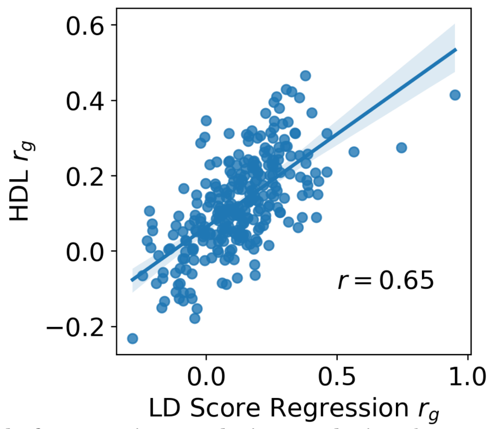


**Fig. A1 Comparison of methods for genetic correlation analysis.** The association between genetic correlation estimates of MIND network edges and FC network edges computed using High-Definition Likelihood (HDL) and LD Score Regression (Ning et al, 2020; Bulik-Sullivan et al, 2015). The mean genetic correlation for both methods was 0.12, and a paired $t$-test demonstrated that the distribution of $r_g$ estimates computed via these two methods were not statistically different ($P = 0.96$)

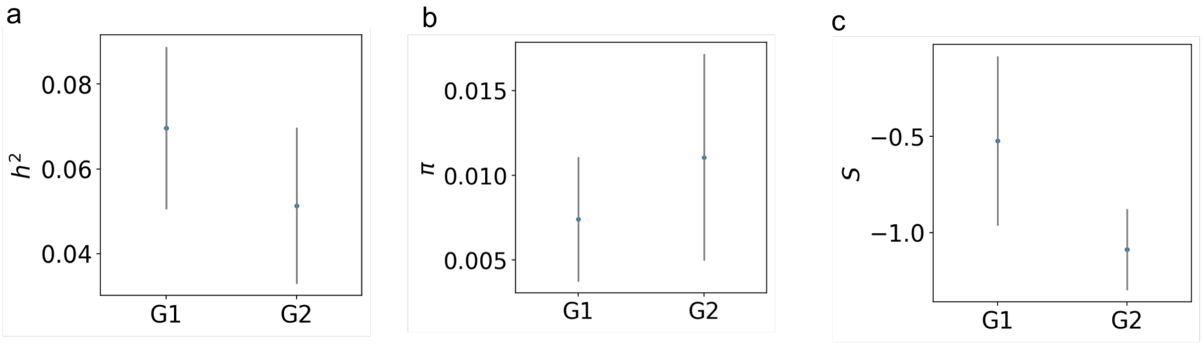


**Fig. A2 Heritability, polygenicity and negative selection of FC along (MIND-derived) G1 and G2.** Estimates of (A) SNP-based heritability, (B) polygenicity parameter $\pi$, and (C) negative selection parameter $S$ of functional connectivity summarised along the two principal MIND genetic gradients. All estimates were calculated using SBayesS (Zeng et al, 2018). Shaded lines indicated 95% confidence intervals.

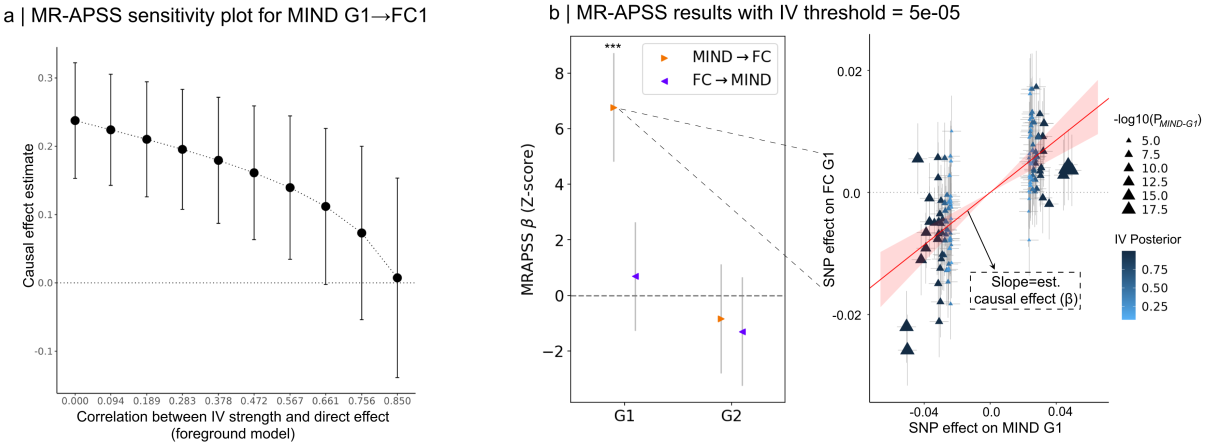


**Fig. A3 Sensitivity analysis of the MR-APSS causal effect of MIND on FC**. A) Sensitivity plot for the result shown in Figure 4E showing a significant causal estimate of MIND G1 on FC1. This plot tests the robustness of the result to model assumptions about the relationship between instrumental variable (IV) strength and the direct causal effect in the foreground model, which is assumed to be zero (Hu et al, 2022). The plot shows here that the positive causal effect of MIND on FC along G1 result remains significant as long as this value remains below 0.661. To put this number in context, the result remains significant for over twice as large a parameter space as the causal evidence for body mass index (BMI) on Type II diabetes, a widely-accepted causal relationship which was previously shown to be robust for values below 0.247 (Hu et al, 2022). B) A replication of the gradient-level MR-APSS results shown in Figure 4E, but with an IV threshold of $5e - 05$ instead of $5e - 04$ . The left plot shows estimated (forward and reverse) causal relationships between MIND and FC (with 95% CI) separately for both G1 and G2. The right plot shows the highly significant causal effect of G1 on FC1, which is shown by the significantly positive slope of the regression plot ($Z$-scored $\beta$=6.2, $P$ = 6.0e-10).

| | Reference | Website | File | $N_{tot}$ | $N_{ca}$ | $N_{co}$ | $N_{eff}$ |
|---|---|---|---|---|---|---|---|
| MDD | Wray et al (2018) | https://pgc.unc.edu/ | daner_pgc_mdd_meta_w2_no23andMe_rmUKBB.gz | 142646 | 45396 | 97250 | 61898 |
| SCZ | Trubetskoy et al (2022) | https://pgc.unc.edu/ | PGC3_SCZ_wave3.european.autosome.public.v3.tsv.gz | 130644 | 53386 | 77258 | 58749 |
| ADHD | Demontis et al (2023) | https://pgc.unc.edu/ | ADHD2022_iPSYCH_deCODE_PGC.meta.gz | 292548 | 38691 | 275986 | 67867 |
| ASD | Grove et al (2019) | https://pgc.unc.edu/ | iPSYCH-PGC_ASD_Nov2017.gz | 46351 | 18,382 | 27,969 | 22184 |
| BPD | Mullins et al (2021) | https://pgc.unc.edu/ | daner_bip_pgc3_nm_noukbiobank.gz | 371549 | 41917 | 371549 | 75334 |
| ALZ | Wightman et al (2021) | https://pgc.unc.edu/ | PGCALZ2ExcludingUKBand23andME_METAL_InverseVariance_MetaAnalysis.txt | 398058 | - | - | 113888 |
| CRP | Said et al (2022) | https://ftp.ebi.ac.uk/pub/databases/gwas/summary_statistics/ | GCST90029070_buildGRCh37.tsv.gz | 575,531 | - | - | - |
| BMI | Yengo et al (2018); Locke et al (2015) | https://portals.broadinstitute.org/collaboration/giant/index.php/GIANT_consortium_data_files#2018_GIANT_and_UK_BioBank_Meta-analysis | Meta-analysis_Locke_et_al+UKBiobank_2018_UPDATED.txt | 795640 | - | - | - |
| GEST | Liu et al (2019) | https://egg-consortium.org/ | Fetal_gest_duration_NComms2019.txt.gz | 84,689 | - | - | - |

**Table A1 External summary statistics accessed and used in this study**. ALZ: Alzheimer's, GEST: gestational age at birth, SCZ: schizophrenia, MDD; major depressive disorder, BMI; body mass index, CRP; C-reactive protein levels, BPD; bipolar disorder, ADHD; attention-deficit hyperactivity disorder, ASD; autism spectrum disorder, $N_{Ca}$; number of cases, $N_{Co}$; number of controls, $N_{Eff}$; Effective sample size, $N_{Tot}$; Total sample size. For MDD, SCZ, ADHD, and ASD, $N_{eff}$ was approximated as $2/(1/N_{Ca} + 1/N_{co})$ as described by Winkler et al (2014), with the maximum $N_{Ca}$ and $N_{Co}$ values over all SNPs used in the calculation. For these traits, $N_{Tot}$ was calculated as these two maximum values summed together. For ALZ, $N_{Ca}$ and $N_{Co}$ were not provided in the summary stats files, so the precomputed maximum $N_{eff}$ and $N_{Tot}$ were used instead. For continuous traits, calculating $N_{eff}$ was not applicable.

| | | IV threshold = 5e-04 | | IV threshold = 5e-05 | | |
|---|---|---|---|---|---|---|
| Exposure | Outcome | $\beta$ | P-val | $\beta$ | P-val | P<0.05 across thresholds |
| G1 | SCZ | 0.042 | 2.0594e-02 | 0.037 | 4.3817e-02 | ✓ |
| G2 | SCZ | -0.13 | 1.6887e-01 | -0.149 | 2.8469e-01 | X |
| G2 | MDD | 0.02 | 4.1479e-01 | 0.0489 | 3.7179e-01 | X |
| G2 | ADHD | -0.1054 | 4.6227e-02 | -0.146 | 1.4359e-01 | X |
| G2 | CRP | -0.1099 | 2.4593e-01 | -0.135 | 3.4173e-01 | X |
| G2 | BMI | -0.0284 | 7.0446e-01 | 0.056 | 3.6625e-01 | X |
| SCZ | G1 | -0.004 | 9.0280e-01 | -0.0018 | 9.6114e-01 | X |
| SCZ | G2 | -0.067 | 9.3939e-02 | -0.078 | 8.4111e-02 | X |
| MDD | G2 | -0.259 | 3.1982e-02 | -0.175 | 2.9114e-01 | X |
| ADHD | G2 | -0.178 | 1.2060e-01 | -0.210 | 1.6264e-01 | X |
| CRP | G2 | -0.081 | 2.6970e-02 | -0.076 | 4.4732e-02 | ✓ |
| BMI | G2 | -0.145 | 5.7496e-04 | -0.151 | 7.9968e-04 | ✓ |

**Table A2 Mendelian Randomization between MIND gradients and biological outcomes.** Mendelian Randomization results for the six significant genetic correlations observed between G1 or G2 and the 9 biological outcomes considered, as indicated by elements of the $r_g$ matrix in Figure 5A (and excluding FC phenotypes). Estimates are derived using MR-APSS using different P-value thresholds for instrumental variable (IV) selection.
